# Neuronal Activity in Orbitofrontal Cortex during Trinary Choices under Risk

**DOI:** 10.1101/2025.09.19.677423

**Authors:** Miguel Barretto-Garcia, Jiaxin Cindy Tu, Camillo Padoa-Schioppa

## Abstract

Economic choice entails computing and comparing the subjective values of different goods. Orbitofrontal cortex (OFC) is thought to contribute to both operations. However, previous work focused almost exclusively on binary choices, raising the question of whether current notions hold for multinary choices. Here we recorded from rhesus monkeys making trinary choices. Offers varied on three dimensions – juice flavor, quantity, and probability. In these experiments, quantity and probability varied continuously within a preset range. Animal choices were generally risk seeking and satisfied independence of irrelevant alternatives (IIA) – a fundamental assumption in standard economic theory. Different neurons encoded the values of individual offers, the choice outcome, and the chosen value – i.e., the same variables previously identified under binary choices. In addition, other cell groups encoded the chosen probability and the chosen hemifield. Notably, the activity of offer value cells reflected the risk attitude and fluctuated from session to session in ways that matched fluctuations observed behaviorally. In other words, the activity of these neurons reflected the subjective nature of value. Importantly, the representation of decision variables in OFC was invariant to changes in menu size – a property that effectively implies IIA.

**SIGNIFICANCE:** Orbitofrontal cortex (OFC) is necessary for the computation and the comparison of subjective values underlying economic choices. However, most previous studies examined choices between two options, and it remains unclear whether current notions apply to multinary choices. Barretto-Garcia and colleagues recorded from the OFC of monkeys choosing between three juice flavors offered in variable quantities and probabilities. Animals’ choices were consistent with the independence of irrelevant alternatives (IIA) – a condition necessary for rational behavior. Different neurons in OFC encoded the values of individual offers, the choice outcome, and the chosen value. The activity of value-encoding cells reflected the animals’ risk attitude. Importantly, the representation of decision variables was invariant to changes in menu size – a property that effectively implies IIA.

## INTRODUCTION

Established literature implicates OFC in economic decisions. Clinical studies in humans and lesion studies in other species indicate that this area is necessary for valuation and choice (Rudebeck et al., 2013; Lak et al., 2014; Reber et al., 2017; Kuwabara et al., 2020; Yu et al., 2022; Gore et al., 2023). Consistent with this view, experiments using electrical stimulation demonstrate that offer values represented in OFC are causal to choices (Ballesta et al., 2020) and that this area also participates in value comparison (Ballesta et al., 2022). Neurophysiology studies identified in OFC neurons encoding individual offer values, the binary choice outcome, and the chosen value (Padoa-Schioppa and Assad, 2006; Pastor-Bernier et al., 2019). Notably, trial-by-trial fluctuations in each cell group correlated with choice variability (Padoa-Schioppa, 2013; McGinty and Lupkin, 2023), and the representation of goods and values was found to be invariant for changes of menu, which implies transitive preferences (Xie and Padoa-Schioppa, 2016). Taken together, these findings provide a rather coherent picture suggesting that OFC hosts a decision circuit (Rich and Wallis, 2016; Padoa-Schioppa and Conen, 2017). However, important questions remain open. Most notably, in many everyday circumstances, individuals choose between multiple options. Yet, the vast majority of previous studies examined choices between two goods (binary choices). A few experiments that examined trinary choices did not focus on OFC (Louie et al., 2013; Chau et al., 2014; Mehta et al., 2019). One study of OFC included trinary choices, but provided only a succinct description of neuronal activity in this condition (Pastor-Bernier et al., 2019). Hence, it remains unclear whether current notions on the neural mechanisms underlying economic decisions generalize to multinary choices. To address this fundamental issue, we recorded from the OFC of monkeys performing a newly designed choice task. In each session, animals chose between three juice flavors offered in variable quantities and probabilities. For each flavor, both quantity and probability varied continuously within fixed ranges. Here we report several results.

First, logistic analyses of choices showed that our animals were generally risk seeking, which confirmed previous findings (Heilbronner and Hayden, 2013). In our experiments, trinary choices and binary choices were randomly interleaved in each session. Importantly, the relative values of the juices, the risk attitude, and the choice accuracy measured in trinary choices were statistically indistinguishable from those measured in binary choices. In other words, monkeys’ choices satisfied IIA – a fundamental assumption in standard economic theory (Kreps, 1990) (see **Discussion**).

Second, we examined neuronal activity in OFC in relation to a large number of candidate variables. As a population, neurons encoded several variables including the offer value of individual juices, binary variables associated with the chosen juice, and the chosen value. These variables capture both the input (offer value) and the output (chosen juice, chosen value) of the decision process, and effectively generalize the variables previously identified for binary choices (Padoa-Schioppa and Assad, 2006).

Third, we investigated the neuronal origins of IIA. Specifically, we analyzed the activity of individual neurons separately in binary and trinary choices, and we compared the results across the population. Individual neurons generally encoded the same variable with the same strength (activity range) in the two sets of trials. In other words, the representation of decision variables in OFC was invariant to changes in menu size. If cell groups identified in OFC constitute the building blocks of a decision circuit, invariance to changes in menu size effectively instantiates IIA.

Fourth, our experimental design provided the opportunity to address a long-standing issue. Economic values are intrinsically subjective, and a fundamental question is whether value-encoding neurons reflect the subjective nature of values. Previous studies demonstrated that neurons encoding the chosen value do. Indeed, their activity was found to vary from session to session in ways that matched fluctuations in the relative value of the offered goods (Padoa-Schioppa and Assad, 2006). In contrast, previous work failed to demonstrate that the activity of offer value cells encodes subjective values, as opposed to some physical property of the offer such as the expected juice quantity (Raghuraman and Padoa-Schioppa, 2014). Here we derived a neuronal measure for the risk attitude from the activity of each offer value cell. This neuronal measure co-varied with, and was statistically indistinguishable from that obtained behaviorally. In other words, the activity of offer value cells indeed reflected the subjective nature of values.

Taken together, our results significantly expand previous notions and deepen our understanding for the neural mechanisms underlying economic decisions.

## METHODS

### Experimental procedures

All experimental procedures conformed to the NIH *Guide for the Care and Use of Laboratory Animals* and were approved by the Institutional Animal Care and Use Committee (IACUC) at Washington University in St Louis.

Two adult male rhesus monkeys (*Macaca mulatta*; monkey E, 10 kg; monkey G, 10 kg) participated in the experiments. After familiarization and chair training, we implanted a head post and an oval recording chamber (main axes 30 mm and 50 mm) under general anesthesia. In both monkeys, the chamber was placed bilaterally, with the longer axis parallel to a coronal plane, and it was centered on 30 mm AP and 0 mm ML. Pre- and post-surgery structural MRIs were used to guide neuronal recordings. During the experiments, monkeys sat in an electrically insulated enclosure (Crist Instruments) in front of a computer monitor (57 cm distance). The gaze direction was monitored with an infrared video camera (Eyelink, SR Research). The behavioral task was controlled through a custom software (Monkeylogic) written in Matlab (MathWorks) (Hwang et al., 2019).

Monkeys performed a trinary choice task (**Fig.1**). In each session, they chose between three juice flavors labeled A, B and C, in (expected) decreasing order of preference. Each juice was offered in variable quantity and probability. Each trial began with the animal fixating a dot at the center of the monitor. After 1 s, three offers appeared. Each offer was represented as an incomplete pie, where the color indicated the juice flavor, the radius indicated the quantity, and the filling angle indicated the probability. The centers of the offers were placed on an invisible circumference 8 cm away from the center fixation point, and the 3 offers were placed 120° from each other in one of two possible configurations (angles 0, 120, 240 or angles 60, 180, 300). The position of each juice flavor (A, B, C) and the spatial configuration varied pseudo-randomly from trial to trial. After a randomly variable delay of 1-2 s, the center fixation point disappeared, and each of the 3 offers was substituted with a colored saccade target (go signal). The animal indicated its choice with a saccade, and had to maintain fixation on the chosen target for an additional 0.75 s. At that point, the trial outcome was revealed – either the animal received the chosen juice, or it heard a “poor luck” sound. The probability, quantity, and flavor of the delivered juice always matched the chosen offer. If the animal broke center fixation prior to the go signal or peripheral fixation prior to juice delivery, the trial was aborted. Otherwise, the trial was termed successful.

**Figure 1.**
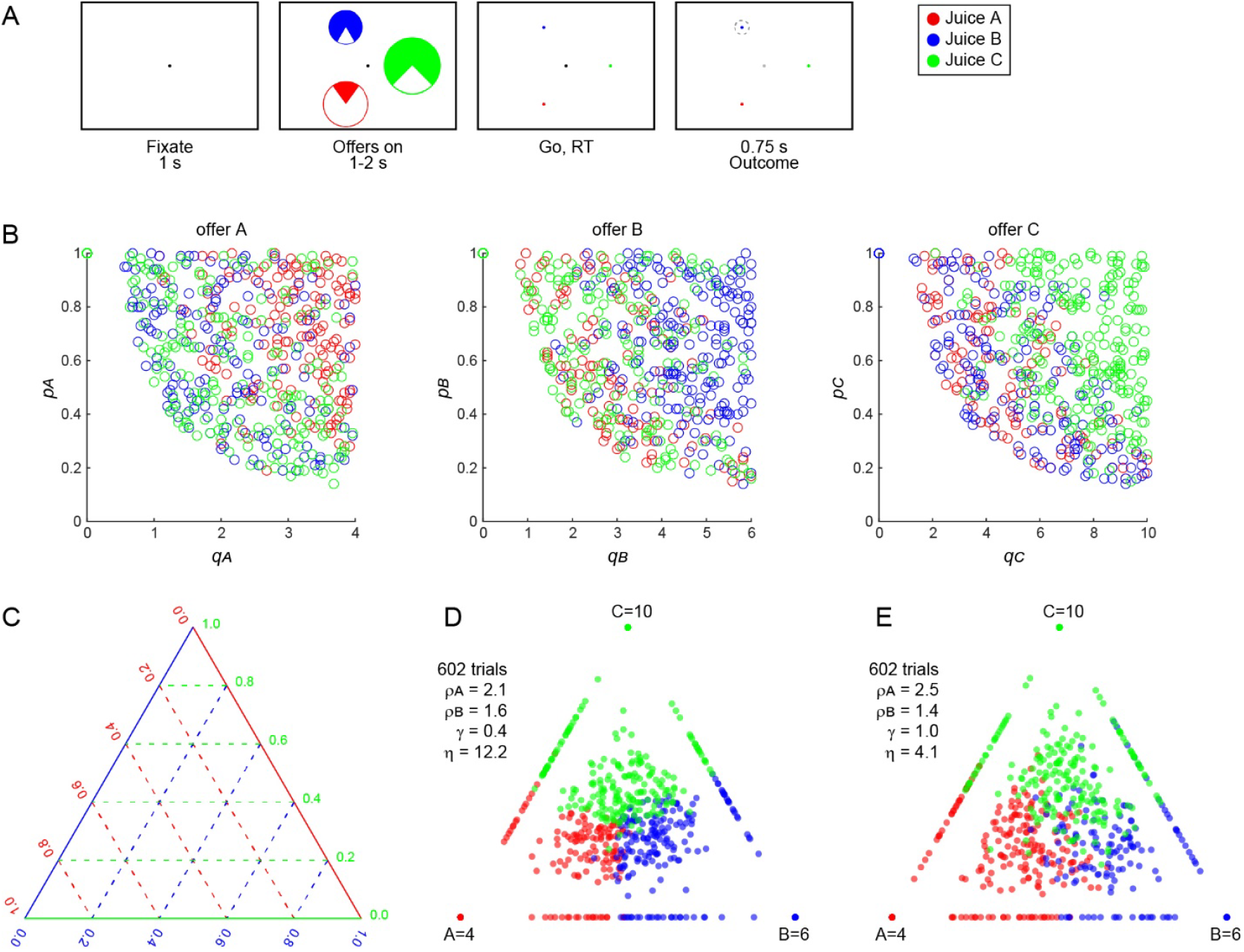
Trinary choice task and simplex representation of choice patterns. **A**. Trinary choice task. Monkeys chose between three juice offers varying in flavor, quantity, and probability. At the beginning of the trial, the animal gazed a fixation point at the center of a computer monitor. After 1 s, three offers appeared on the monitor. Each offer was depicted as an incomplete pie, with the color indicating juice flavor, the radius indicating the quantity, and the filled angle indicating the probability. The centers of the three pie images were arranged on a circle around the center fixation point (at 8.0° distance) and were spaced 120° from each other in one of two symmetric configurations (angles 0°, 120°, 240° or 60°, 180°, 300°). The animal maintained center fixation for a randomly variable delay (1-2 s), at the end of which the center fixation point was extinguished and the three offers were substituted by three saccade targets (go signal). The animal indicated its choice with a saccade and maintained peripheral fixation for an additional 0.75 s. At the end of this delay, we delivered the chosen juice *J* in quantity *q*_*J*_ with probability *p*_*J*_; with probability 1 − *p*_*J*_, we did not deliver any juice and played a “poor luck” sound. **B**. Joint distribution of quantities and probabilities, example session. The three panels illustrate the distribution of offers for juices A, B, and C. In each panel, the x-axis represents the offer quantity, the y-axis represents the offer probability, and each data point is one trial. Colors indicate the chosen juice. Offers were uniformly distributed in the plane (p, q), with the constraint that the expected quantity be ≥ a fixed minimum. **C**. Simplex representation of 3D data in 2D. Here, each dimension represents the subjective value *V*_*J*_ of one juice offer (J = A, B, C). *V*_*J*_ depends on the relative value of juice J (*ρ*_*J*_) and on the risk attitude (*γ*; **Eq.24**). For the simplex plot, each subjective value was divided by the maximum available for that juice type in the session (i.e., we computed *v*_*J*_ = *V*_*J*_⁄*Q*_*J*_, where *Q*_*J*_ is the maximum possible *q*_*J*_ for the session). Then, the three values were normalized to barycentric coordinates (i.e., we computed *x*_*J*_ = *v*_*J*_⁄(*v*_*A*_ + *v*_*B*_ + *v*_*C*_)). The position of each data point in the simplex was determined by *x*_*A*_, *x*_*B*_, and *x*_*C*_. Dashed lines in panel C indicate sets of points for which one of the *x*_*J*_ is fixed. For example, proceeding from the bottom side of the triangle to the top vertex, *x*_*C*_ increases from 0 to 1 (green dashed lines). **D**. Choice pattern, example session 1. Each data point represents one trial. The position of the data point is determined by the three offer values (per panel C) and the color indicates the animal’s choice. Data points on the borders of the triangle correspond to binary choices; data points on the vertices of the triangle correspond to forced choices. This session included 602 trials and we set *Q*_*A*_ = 4, *Q*_*B*_ = 6, and *Q*_*C*_ = 10. Relative values (*ρ*_*A*_, *ρ*_*B*_), risk attitude (*γ*), and choice accuracy (*η*) derived from the logistic analysis are indicated in the panel. Notably, the three clouds of data points corresponding to the three chosen juices are well separated (high *η*). **E**. Choice pattern, example session 2 (602 trials). Same format as in panel D. In this case, the three clouds of data points corresponding to the three chosen juices are more overlapping (low *η*).

In most trials, 3 juices were offered (60% trinary choices; A:B:C trials). In a fraction of trials only 2 juices were offered (30% binary choices; A:B, B:C, C:A trials in equal proportions). In a smaller fraction of trials, only 1 juice was offered (10% forced choices; A, B, C trials in equal proportions). For each session and for each juice flavor, offer quantity and offer probability varied continuously within a preset range. The maximum quantity differed between juices and was set at the beginning of each session based on the anticipated relative values, in such a way that the animal would choose each flavor in roughly equal fraction of trials. In each trial and for each juice *J, p*_*J*_ and *q*_*J*_ were selected randomly from a joint uniform distribution, imposing the constraint that the expected quantity be not too small (*p*_*J*_*q*_*J*_ ≥ *Q*/8). Sessions typically lasted 170-725 trials. Across sessions and monkeys, we used 11 juice types and 36 different combinations of juices. The juice quantum – i.e., the volume of juice corresponding to 1 cm of pie radius – was set at 90-100 μl in different sessions, and was fixed within each session.

Neuronal recordings focused on area 13m in the central orbital gyrus (Ongur and Price, 2000) and were guided by MRI scans. We recorded from both hemispheres of monkey E (left: AP 31:35, ML – 8:–10; right: AP 31:35, ML 6:10) and both hemispheres of monkey G (left: AP 31:36, ML –7:–12; right: AP 31:36, ML 4:9). Extra-cellular recordings were conducted using tungsten electrodes (125 µm shank diameter; FHC) or V-probes (16 channels; Plexon). Electrodes or probes were advanced remotely using a custom-built motorized micro-drive (step size 2.5 µm). Typically, one motor advanced two electrodes placed 1 mm apart, and 1-2 such pairs of electrodes were advanced unilaterally or bilaterally in each session. Alternatively, we recorded from a single V-probe. Signals were amplified (gain: 10,000), filtered (high-pass cutoff: 300 Hz; low-pass cutoff: 6 kHz; Lynx 8, Neuralynx), digitized (frequency: 40 kHz), and saved to disk (Power 1401, Cambridge Electronic Design). Spike sorting was performed off-line (Spike 2 v6, Cambridge Electronic Design). Only cells that appeared well isolated and stable throughout the session were included in the analysis. All subsequent analyses were performed in Matlab (version R2022b; MathWorks).

### Logistic analysis of choice data

Behavioral choice data recorded in each session were submitted to logistic analysis (Padoa-Schioppa, 2022). In the log value ratio logistic model, the probability of choosing juice *J* is:

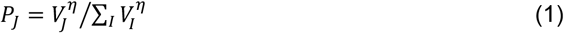

where *V*_*J*_ is the value of offer *J, I* = A, B, C, and *η* is the choice accuracy (also termed inverse temperature). For juice offers that vary in flavor, quantity and probability, the offer value of juice *J* is:

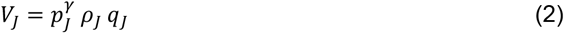

where *q*_*J*_ and *p*_*J*_ are the quantity and probability associated with offer *J*, and *ρ*_*J*_ is the subjective value of a unit volume of juice *J*. Without losing generality, we express subjective values in units of juice C. In these units, *ρ*_*C*_ = 1, and *ρ*_*A*_ and *ρ*_*B*_ are termed relative values of juices A and B, respectively. In **Eq.2**, *γ* quantifies how subjective probabilities are distorted compared to actual probabilities. Specifically, *γ* < 1 means that the monkey treats probabilities as higher than they are (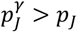 risk seeking); *γ* > 1 means that the monkey treats probabilities as lower than they are (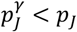 risk aversion); *γ* = 1 means that the monkey treats the probabilities exactly as they are (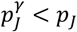 risk indifference). The last condition yields maximum expected payoff. We refer to *γ* as the risk attitude.

Substituting **Eq.2** in **Eq.1**, the probability of choosing each of the three offers can be written explicitly:

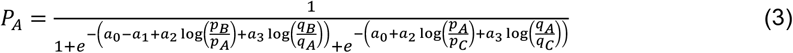

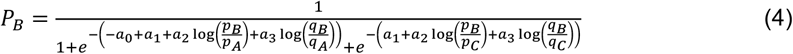

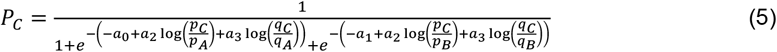

In **Eqs.3-5**, we defined parameters *a*_0_ = *η* log(*ρ*_*A*_), *a*_1_ = *η* log(*ρ*_*B*_), *a*_2_ = *η γ*, and *a*_3_ = *η*.

For each session, we estimated regression parameters *a*_0_ … *a*_3_ with maximum likelihood, using the objective function:

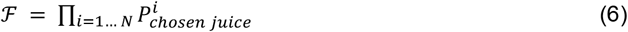

where *N* was the number of trials in the session and 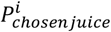 was the probability that on trial *i* the animal would choose the juice it actually did choose (**Eqs.3-5**). From fitted parameters 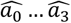, we computed the values of behavioral parameters 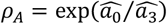, 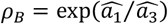, 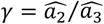, and 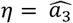 (elsewhere, hats 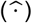 have been omitted for readability).

### Testing the independence of irrelevant alternatives (IIA)

To assess whether choices violated IIA, we performed the logistic analysis separately in binary choice trials and in trinary choice trials, and we compared the two sets of parameters. This approach is similar to that of the Hausman-McFadden test (Hausman and D., 1984; Train, 2003), which effectively simulates binary choices, except that binary choices were actually included in our data set. Note, however, that our experimental design allowed a more powerful test, because binary choices were actually included in each session, and not simulated as counterfactual. In practice, the logistic analysis described above provided two sets of parameters 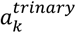 and 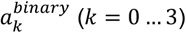, from which we computed the behavioral parameters 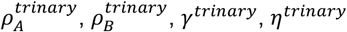 and 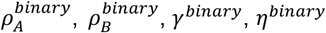. Each behavioral parameter was compared across sessions using a paired, two-sided t-test; in particular, we examined whether the choice variability differed *η* between binary and trinary choices.

In addition, to perform a comprehensive comparison of the fitted parameters 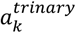 and 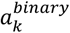, we derived for each parameter *a*_k_ the statistics

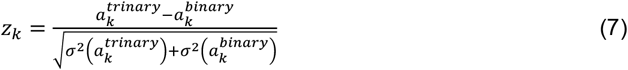

where 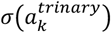 and 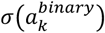 were standard errors obtained from the logistic regression. We then computed the corresponding *p*_*k*_ as a complementary error function

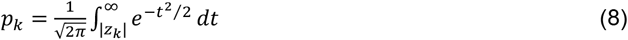

Each *p*_*k*_ is essentially a p value – i.e., parameter *a*_*k*_ differs significantly between trinary and binary choices if *p*_*k*_ < 0.05. Finally, we combined all the *p*_*k*_ and computed an overall measurement using Fisher’s method, *F*,

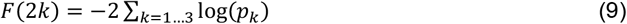

*F* follows a *X*^2^ distribution with 2*k* degrees of freedom. A *X*^2^ test provided a comprehensive p value for the session, namely *p*_*Fisher*_.

### Neuronal activity profiles

Several figures illustrate the activity profiles of individual cells. To calculate the activity profiles, we first divided trials according to the relevant variable. Trials were then aligned at the offer onset and separately at the trial outcome. For each alignment and each trial, the spike train was smoothed using the method used in previous studies (So and Stuphorn, 2010; Padoa-Schioppa, 2013). Spike times, expressed in 1 ms resolution, were convolved with the kernel:

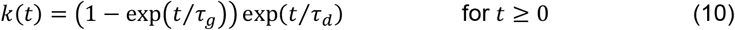

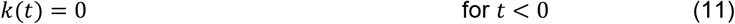

This kernel mimics a post-synaptic potential and ensures that spikes exert influence only forward in time. Specifically, we used *τ*_*g*_ = 1 ms and *τ*_*d*_ = 20 ms. For each cell and for each group of trials, we averaged spike trains across trials and we obtained a smoothed activity profile. For display purposes, we coarse-grained the resulting signal by averaging the activity in 5 ms bins (non-overlapping). Activity profiles were only computed for illustration purposes; all statistical analyses were conducted on spike counts.

### Variable selection analysis

Neuronal data were analyzed in 8 time windows: pre-offer (0.5 s preceding the offer onset), post-offer (0.5 s after the offer onset), late delay (0.5–1.0 s after the offer onset), pre-go (0.5 s preceding the go cue), reaction time (from the go cue to the saccade start), pre-juice (0.5 s preceding juice delivery), post-outcome (0.5 s after juice delivery), and post-outcome2 (0.5–1.0 s after juice delivery). We refer to the activity of one cell in one time window as a neuronal response. Our primary goal was to assess what decision variables are represented in OFC during trinary choices. Following previous work, we defined a series of variables that OFC neurons could conceivably encode. These included variables associated with a single juice (*quantity of A, probability of A, expected quantity of A, offer value A, quantity of B*, etc.), a spatial variable identifying the *chosen hemifield*, variables associated with the chosen offer (*A chosen, B chosen, C chosen, chosen quantity, chosen probability*, etc.), and variables reflecting the trial outcome – i.e., whether juice was eventually delivered or whether it was a poor luck trial (*got juice, received value*, etc.). In total, we examined 29 variables (**Table 1**).

**Table 1.**
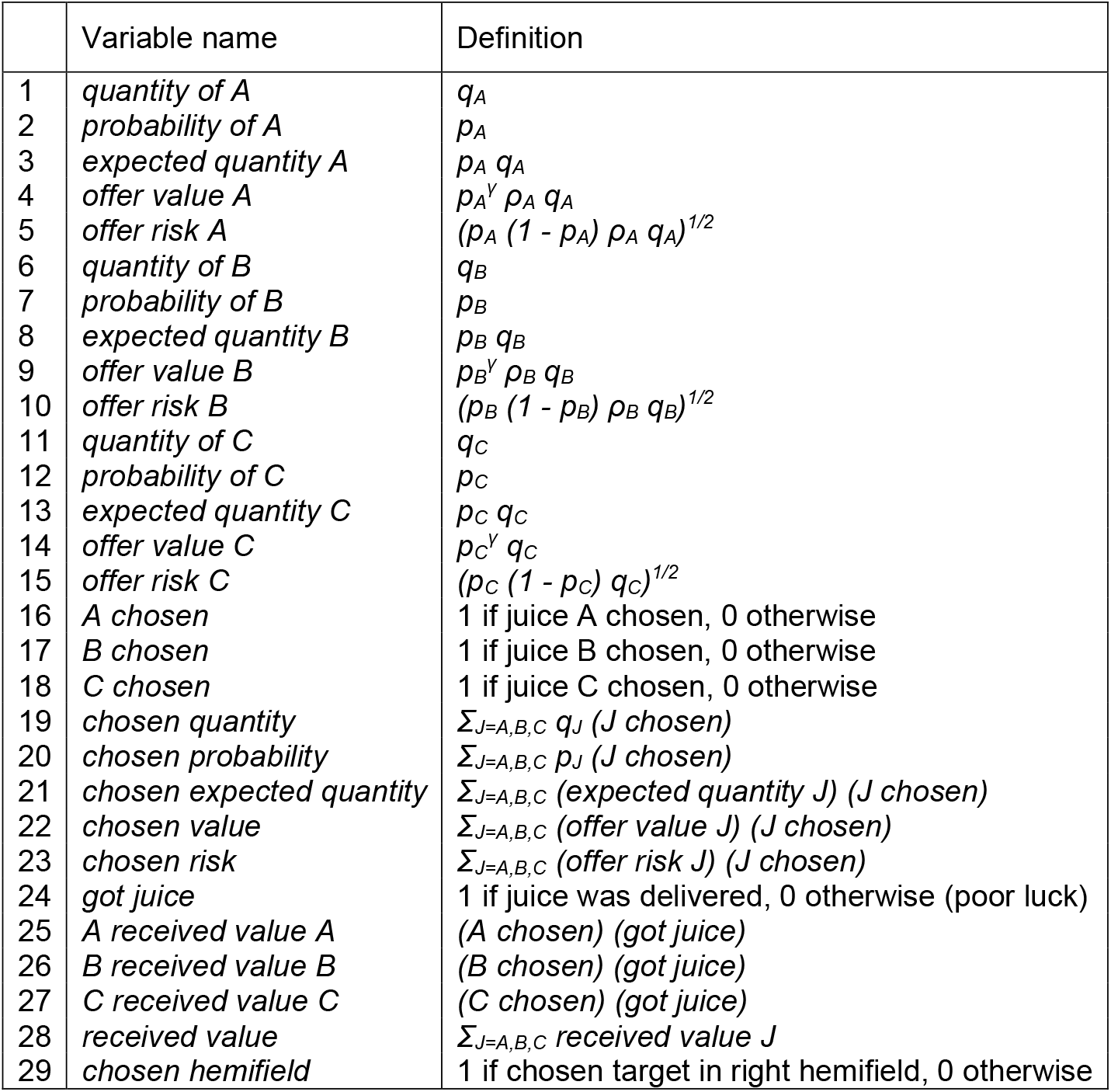
Task variables examined in this study.

Ultimately, we wanted to identify a small subset of variables that would explain most of the neuronal activity. Compared to previous studies (Padoa-Schioppa and Assad, 2006; Raghuraman and Padoa-Schioppa, 2014), a key difference in our experimental design was that offered quantities and probabilities varied continuously and were not quantized. Thus we could not define trial types or conduct an ANOVA to identify task-related cells. One possible approach was to regress single-trial firing rates against each of the variables under consideration. However, standard linear regressions assume that data points come from Gaussian distributions of equal variance, and spike counts violate both these assumptions (Conen and Padoa-Schioppa, 2015). This issue is greatly mitigated by regressing firing rates averaged across trials. Hence, for each variable, we divided trials into 20 quantiles, each including the same number of trials. Indicating with 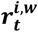 the firing rate of neuron ***i*** recorded in time window ***w*** and trial ***t***, for each quantile ***q***, we computed the mean firing rate across trials 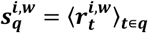, where ⟨⋅⟩ indicates the average. We refer to 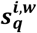 as binned firing rates. We then regressed binned firing rates against the variable under examination, and we obtained the regression slope and the R^2^. If the regression slope differed significantly from zero (p<0.01), the variable was said to explain the response. If the variable did not explain the response, we set R^2^ = 0. We repeated this operation for each variable. Notably, the binning was done separately for each variable such that trials binned together for one particular variable could be binned separately for other variables. Any given response could be explained by multiple variables. In this case, we identified the variable providing the best explanation (highest R^2^). Neurons whose activity was explained by ≥1 variable in ≥1 time window were termed task related; neurons that were not explained by any variable in any time window were referred to as *untuned*.

From here on, the variable selection analysis proceeded as in previous studies (Padoa-Schioppa and Assad, 2006). To identify a small number of variables that best accounted for the whole population, we used a best subset procedure. For *k* = 1,2,3… we examined all the subsets of *k* variables, we computed the total R^2^ explained by each subset, and we identified the subset that explained the maximum total R^2^. We repeated this procedure until when the best subset explained at least 70% of the total R^2^ explained by the full set of 29 variables. Importantly, the variable selection analysis was conducted imposing a consistency constraint across juice flavors: for any value of *k*, we only examined subsets that included all three analogous variables (e.g., *offer quantity A, offer quantity B, offer quantity C*). This constraint ensured that results of the analysis would be interpretable.

### Neuronal origins of IIA

We aimed to assess whether the neuronal representation in OFC is invariant to changes in menu size – i.e., whether the encoding of decision variables during trinary choices differs from that during binary choices. To address this issue, we focused on pre-outcome time windows and on task-related responses. We conducted two analyses.

First, for each neuronal response, we separated binary and trinary choice trials (including forced choices in both sets). We then analyzed each set of trials and assigned the response to one of the variables identified in the variable selection analysis based on the R^2^. We then pooled responses and we constructed a 10×10 contingency table where rows and columns were the decision variables in binary and trinary choice trials, respectively, and entries were cell counts. We refer to entry (*i, j*) in this matrix as *X*_*i*,*j*_. To assess whether cell counts deviated from chance, we computed the corresponding table of odds ratios (ORs), For each entry (*i*,*j*), OR was defined as follows:

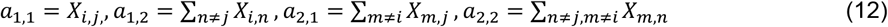

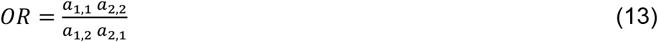

The table of ORs quantifies the association strength between the two classifications. For each entry in the table, OR < 1, OR = 1, or OR > 1 mean that the corresponding cell count was lower than, equal to, or larger than expected by chance. Departures from chance were tested with two-tailed Fisher’s exact test.

As a reminder, in each session, roughly 60%, 30%, and 10% or trials were trinary choices, binary, and forced choices, respectively. Because binary and trinary choices differed in number of trials roughly by a 1:2 ratio, the classification in binary choices was necessarily noisier, and thus we expected to observe some classification discrepancy across conditions. To evaluate the effect of the difference in number of trials, we conducted a bootstrap analysis. For each session, we indicate with *n* and *N – n* the number of binary choice trials and the number of trinary choice trials, respectively. For each neuronal response, we pooled binary and trinary choices, we divided them randomly and arbitrarily in two sets of *n* and *N – n* trials, and we repeated the classification analysis described above. We repeated this operation 100 times, we pooled neuronal responses, and we constructed a 10×10 contingency table where rows and columns were the decision variables encoded in when the number of trials was small (*n*) and when the number of trials was large (*N – n*), respectively. We then constructed the corresponding table of ORs. Finally, we compared the contingency table obtained for binary and trinary choices with that obtained for small and large number of trials using a Breslow-Day test (Agresti, 2019).

Second, we examined whether the strength of the encoding – i.e., the neuronal activity range (AR) – differed between binary and trinary choices. Again, for each response, we separated binary and trinary choice trials. For each set of trials, we regressed firing rates against the encoded variables (with the same binning procedure described above). In other words, we performed the linear regression

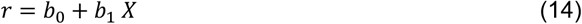

Where *X* was the encoded variable and *b*_0_ and *b*_1_ are the intercept and slopes, respectively. We computed the maximum (*R*_*max*_) and minimum (*R*_*min*_) firing rates,

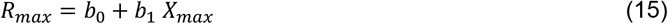

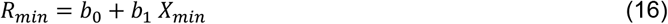

where *X*_*max*_ and *X*_*min*_ are the maximum and minimum value range of the encoded variable, *X*. We then computed the activity range, AR

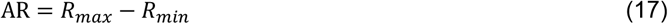

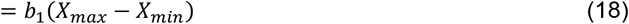

We then computed the difference in activity ranges ΔAR = AR_trinary_ – AR_binary_. We then examined the distribution of ΔAR across the population of neuronal responses, separately for each encoded variable.

Since binary choice trials were roughly half of trinary choices trials, we expected to observe some difference across conditions (indeed, we found that mean(ΔAR) < 0 for all variables). To evaluate the effect of this factor, we conducted a bootstrap analysis. For each neuronal response, we pooled binary and trinary choices, we divided them arbitrarily in two sets of *n* and *N – n* trials, and we repeated the analysis of ARs. We repeated this operation 100 times, and we examined the ΔAR across the population for each variable. Finally, for each variable, we compared the distribution obtained for binary and trinary choices with that obtained for small and large number of trials using a two-sample t-test.

### Neuronal measures of risk attitude

We aimed to assess whether value-encoding cells in OFC reflected the subjective risk attitude *γ*. To do so, we developed a method to derive a measure for the risk attitude (*γ*_*neuron*_) from each neuronal response. This measure was then compared with that obtained from the logistic analysis (*γ*_*behavior*_) in a population analysis. The method used to derive *γ*_*neuron*_ generalized to the case of continuous variables a method previously adopted for quantized variables (Raghuraman and Padoa-Schioppa, 2014). To simplify the notation, in this section we omit the juice subscript *J*.

First, we considered responses encoding the offer value. We examined firing rates in relation to probability *p* and quantity *q*. For each response, we considered the distribution of *p*, according to which we divided trials into 6 quantiles. Each probability quantile included the same number of trials. We indicated with *p*_*k*_ the mean *p* within quantile *k* (*k* = 1…6). For each probability quantile *k*, we regressed the firing rate *r* against the quantity *q*

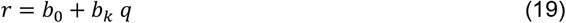

Note that the intercept *b*_0_ – i.e., the baseline firing rate – was the same for all quantiles, while the slope *b*_*k*_ was computed separately for each quantile. The linear regression in **Eq.19** was performed using the binning procedure described above and dividing *q* in 10 bins.

That a neuronal response encodes the *offer value* means that *b*_*k*_ *q* is proportional to the subjective value of the offered juice, which is expressed in **Eq.2**. That is:

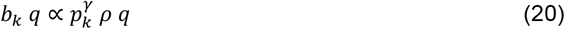

where ∝ indicates proportionality. Taking the log of **Eq.20**, we obtain the relation

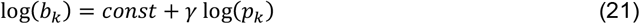

Thus a linear regression of log(*b*_*k*_) onto log(*p*_*k*_) provided a measure for *γ*_*neuron*_.

We proceeded in similar ways for responses encoding the chosen value. For each response, we divided the distribution of probabilities *p* in 6 quantiles, each with the same number of trials. For each quantile *k* = 1…6, we regressed firing rates against values *ρ q*.

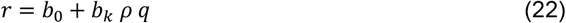

In **Eq.22**, *ρ* and *q* were the relative value and quantity of the chosen juice. Combining **Eq.22** and **Eq.2** and taking the log, we obtained a prediction formally identical to **Eq.21**. Thus a linear regression of log(*b*_*k*_) onto log(*p*_*k*_) provided a measure for *γ*_*neuron*_. For negatively tuned responses, we used |*b*_*k*_ | (i.e., rectified slopes).

These analyses were restricted to early time windows preceding the trial outcome.

## RESULTS

### Choice task and logistic analysis of trinary choices

Two rhesus monkeys, E and G, participated in the experiments. In each session, we used three juice flavors labeled A, B, and C in decreasing order of preference. In 60% of trials, the animal chose between three offers presented simultaneously (trinary choices). In 30% of trials, only two offers were presented (binary choices). In the remaining 10% of trials, only one offer was presented (forced choices). Offers varied on three dimensions – juice flavor, quantity, and probability. Thus, each trial comprised three offers, one per flavor. Each offer was represented visually by an incomplete pie, with the color indicating the juice flavor (*J*), the radius indicating the juice quantity (*q*_*J*_), and the filled angle indicating the probability (*p*_*J*_) (**Fig.1A**). Across trials, *p*_*J*_ and *q*_*J*_ varied continuously over ranges set at the beginning of each session. Specifically, maximum quantities *Q*_*J*_ were set based on the relative values (see below) foreseen for the three juices, and such that the animal would choose each flavor roughly 1/3 of the time; probabilities always varied between 0 and 1. Importantly, we imposed that the expected quantity of each offer be always at least 12.5% of the maximum offer range – in formulas, *p*_*J*_ *q*_*J*_ ≥ *Q*_*J*_⁄8 (**Fig.1B**). Trinary, binary, and forced choices were randomly interleaved. Sessions included 170-725 successful trials (median = 601 trials from monkey E, 431 trials from monkey G). The full dataset included 248 sessions (121 from monkey E, 127 from monkey G).

In each session, choice data were submitted to a logistic analysis based on the following log value ratio model (see **Methods**):

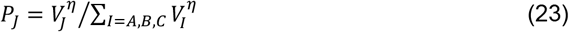

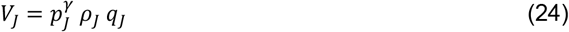

where *J* = A, B, C; *P*_*J*_ is the probability that in the trial of interest the animal chose juice *J*; *V*_*J*_ is the subjective value of offer *J*; *p*_*J*_ and *q*_*J*_ are the probability and quantity of juice *J*, respectively; and *ρ*_*J*_, *γ*, and *η* are parameters characterizing the behavior of the monkey. Specifically, *ρ*_*J*_ captures the subjective value of one unit volume of juice *J*. We conventionally express values in units of juice C, such that *ρ*_*C*_ = 1. Parameters *ρ*_*A*_ and *ρ*_*B*_ are termed relative values of juices A and B, respectively, and were estimated separately for each session. Parameter *γ* is termed risk attitude and captures the subjective distortion of probabilities; conditions *γ* < 1 (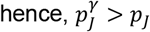), *γ* = 1 (hence, 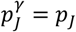), and *γ* > 1 (hence, 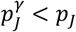) correspond to risk seeking, risk neutrality, and risk aversion, respectively. Finally, parameter *η* is termed choice accuracy, and captures the degree of consistency in the animal’s choices. Intuition about this point is gained by noting that, given three offer values, the probability of choosing the highest value increases as *η* becomes larger (**Eq.23**); conversely, as *η* approaches 0, choices become random.

In these experiments, visualizing fully specified offers would require 6 dimensions (3 quantities, 3 probabilities) and is not feasible. To depict the behavior in one session, one possibility is to illustrate offers in coordinates defined by expected quantities (*p q*) or subjective offer values (*p*^*γ*^*ρ q*), but even in three dimensions it is hard to render the data structure. Thus we used simplex plots, which display 3D data in 2D following normalization and using barycentric coordinates (**Fig.1C**). Axes in the simplex plot were based on the subjective offer values, and each data point was assigned a color according to the chosen juice. **Fig.1DE** illustrate the behavior recorded in 2 example sessions with different levels of risk attitude and choice accuracy.

### Monkeys’ choices satisfy IIA

IIA is a cornerstone concept in standard economic theory. Consider a subject choosing between two options X and Y or, alternatively, between three options X, Y, and Z. Indicating with *P*_*X*_ and *P*_*Y*_ the likelihood of choosing options X and Y, respectively, IIA is satisfied if the ratio *P*_*X*_⁄*P*_*Y*_ does not depend on whether option Z is offered. Conversely, if the presence of option Z alters the ratio *P*_*X*_⁄*P*_*Y*_, IIA is violated. Previous behavioral work indicated that choices in humans and other species can violate IIA in a variety of ways. However, these effects have been found to depend on the experimental design (see **Discussion**). Thus we sought to assess whether our monkeys’ choices were consistent with IIA.

To do so, we repeated the logistic analysis described above (**Eq.23, Eq.24**) restricting it exclusively to binary choices and, separately, exclusively to trinary choices. We thus obtained two sets of parameters 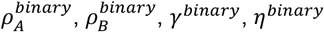 and 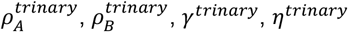. In essence, IIA implies that the two sets of parameters are indistinguishable from each other; conversely, IIA is violated if the parameters obtained for binary and trinary choices differ significantly from each other. We examined this issue in two ways. First, we conducted a population analysis across sessions. In principle, any parameter could vary between binary and trinary choices. In particular, since trinary choices presumably require more cognitive resources, a plausible prediction was that choice accuracy would be lower for trinary than for binary choices (*η*^*trinary*^ < *η*^*binary*^). In fact, our analyses did not find any systematic differences between the parameters measured in the two groups of trials (**Fig.2A-D**). In particular, we did not find significant differences between *η*^*binary*^ and *η*^*trinary*^ (*p* = 0.11; paired, two-sided t-test; **Fig.2E**).

**Figure 2.**
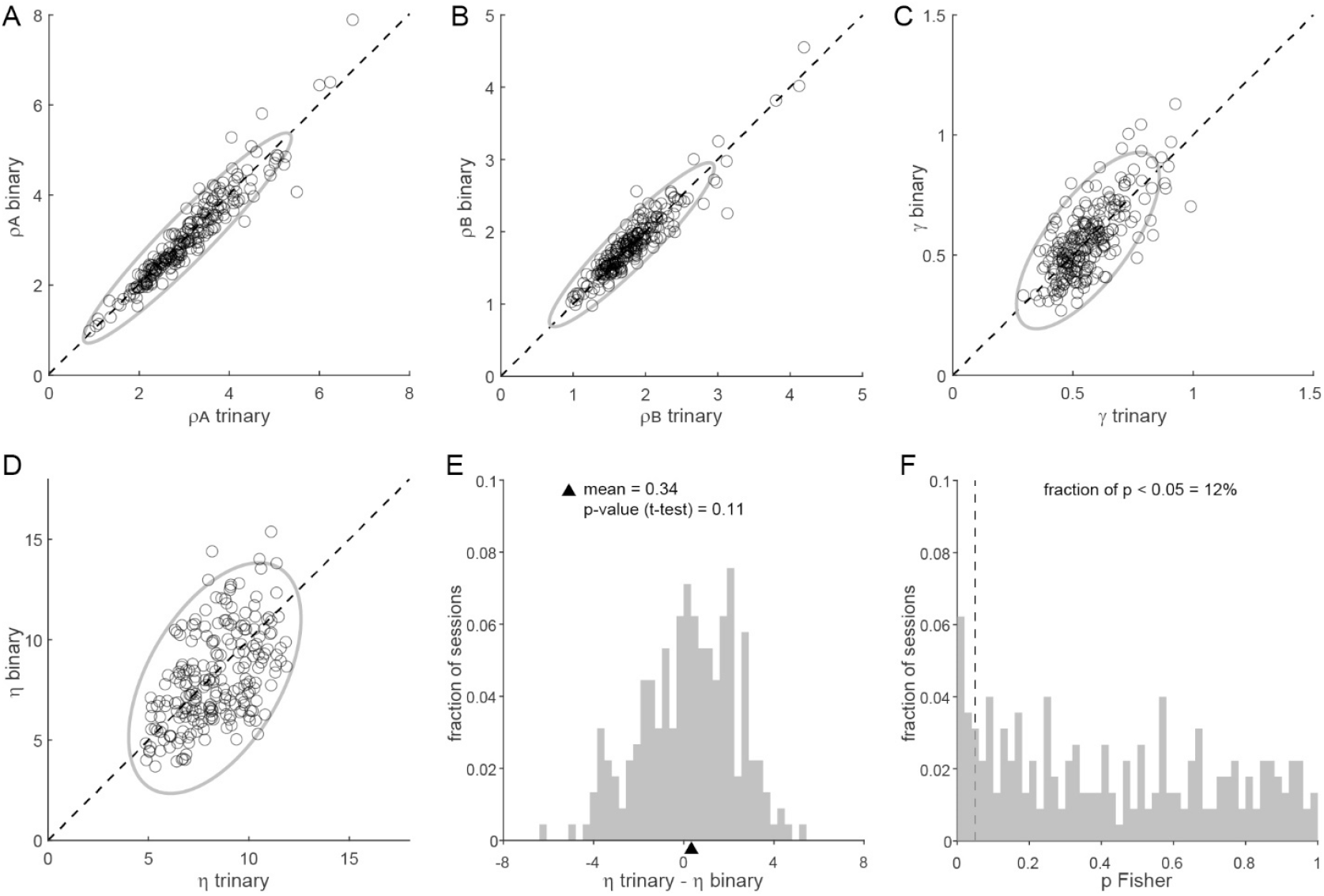
Monkeys choices satisfy independence of irrelevant alternatives (IIA). **AB**. Relative values measured in binary and trinary choices are indistinguishable. Logistic analysis was conducted separately on binary and trinary choice trials. In panel A, x- and y-axis represent the relative value *ρ*_*A*_ measured in the two sets of trials, and each data point represents one session. The dashed line indicates identity and the ellipse indicates 95% confidence interval for the distribution of data points. The two measures of *ρ*_*A*_ did not present any systematic difference (p = 0.231, paired two-sided t-test). Similar conclusions hold for *ρ*_*B*_ (p = 0.384 paired t test; panel B). **C**. Monkeys have the same risk attitude in binary and trinary choices (p = 0.795, paired t test). **DE**. Binary and trinary choices are equally accurate. Panel D has the same format as in panel A. In general, measures of choice accuracy (*η*) obtained from binary choices (y-axis) were affected by larger errors compared to those obtained from trinary choices (x-axis). This is because the number of trials for binary choices was roughly half of that for trinary choices. However, at the population level, the two measures did not present any systematic difference (p = 0.114, paired t test). **F**. Combined assessment of IIA violations. Fisher’s method assessed whether the raw coefficients obtained from the logistic analysis of binary and trinary choices were statistically different. The histogram represents the distribution of p values obtained from Fisher’s test across sessions (see **Methods**). The vertical dashed line indicates the p = 0.05 threshold. For sessions to the right of the dashed line, Fisher’s test failed to detect significant violations of IIA. N = 248 sessions were included in these analyses.

Second, in a complementary analysis, we used Fisher’s method to derive a comprehensive statistic *F* that compared all four parameters obtained from the logistic analysis of binary versus trinary choices (see **Methods**). This statistic followed a *X*^2^ distribution with 8 degrees of freedom. Thus a *X*^2^ test provided a comprehensive p value for the session, namely *p*_*Fisher*_. The distribution of *p*_*Fisher*_ across the whole data set revealed that our monkeys’ choices violated IIA only in a small fraction of sessions (**Fig.2F**).

Importantly, these results held true for individual animals. In particular, *η*^*binary*^ and *η*^*trinary*^ were statistically indistinguishable in both monkey E and monkey G (*p* = 0.40 and *p* = 0.16, respectively; paired, two-sided t-test). Thus we concluded that choices in our experiments were generally consistent with IIA. Of note, inspection of **Fig.2D** also revealed that our monkeys were generally risk seeking (*γ* < 1) in both binary and trinary choices – a result that confirmed previous findings (Heilbronner and Hayden, 2013; Raghuraman and Padoa-Schioppa, 2014).

### Neuronal representation of decision variables

While monkeys performed the trinary choice task, we recorded the activity of 1,469 neurons in central OFC (921 cells from monkey E; 548 cells from monkey G). We conducted a series of analyses to identify what decision variables might be represented in OFC. Notably, our task design was considerably more complex than that of previous studies (Padoa-Schioppa and Assad, 2006; Raghuraman and Padoa-Schioppa, 2014) because animals were offered 3 options, because each option varied on 3 dimensions, and because offer quantities and probabilities varied on a continuum (i.e., they were not quantized). One implication is that trials could not be naturally grouped into a small number of discrete trial types. However, trials could be binned by quantiles of specific variables. With these premises, we first examined the activity profiles of individual cells.

**Fig.3A-G** illustrates the activity of one example neuron in relation to the three offer values. In panel B, trials were divided into 5 quantiles according to the variable *offer value A* (each quantile had the same number of trials). The cell activity appeared strongly modulated by this variable throughout the trial. Indeed, the firing rate recorded in the 500 ms following the offer onset (post-offer time window) was in close linear relation with the *offer value A* (**Fig.3C**). In contrast, the activity of the same neuron was not detectably modulated by either the *offer value B* or the *offer value C* (**Fig.3D-G**). **Fig.3H-N** illustrates another example neuron. In this case, firing rates during offer presentation (late delay time window) were linearly related to the *offer value C* and independent of *offer value A* and *offer value B*.

**Figure 3.**
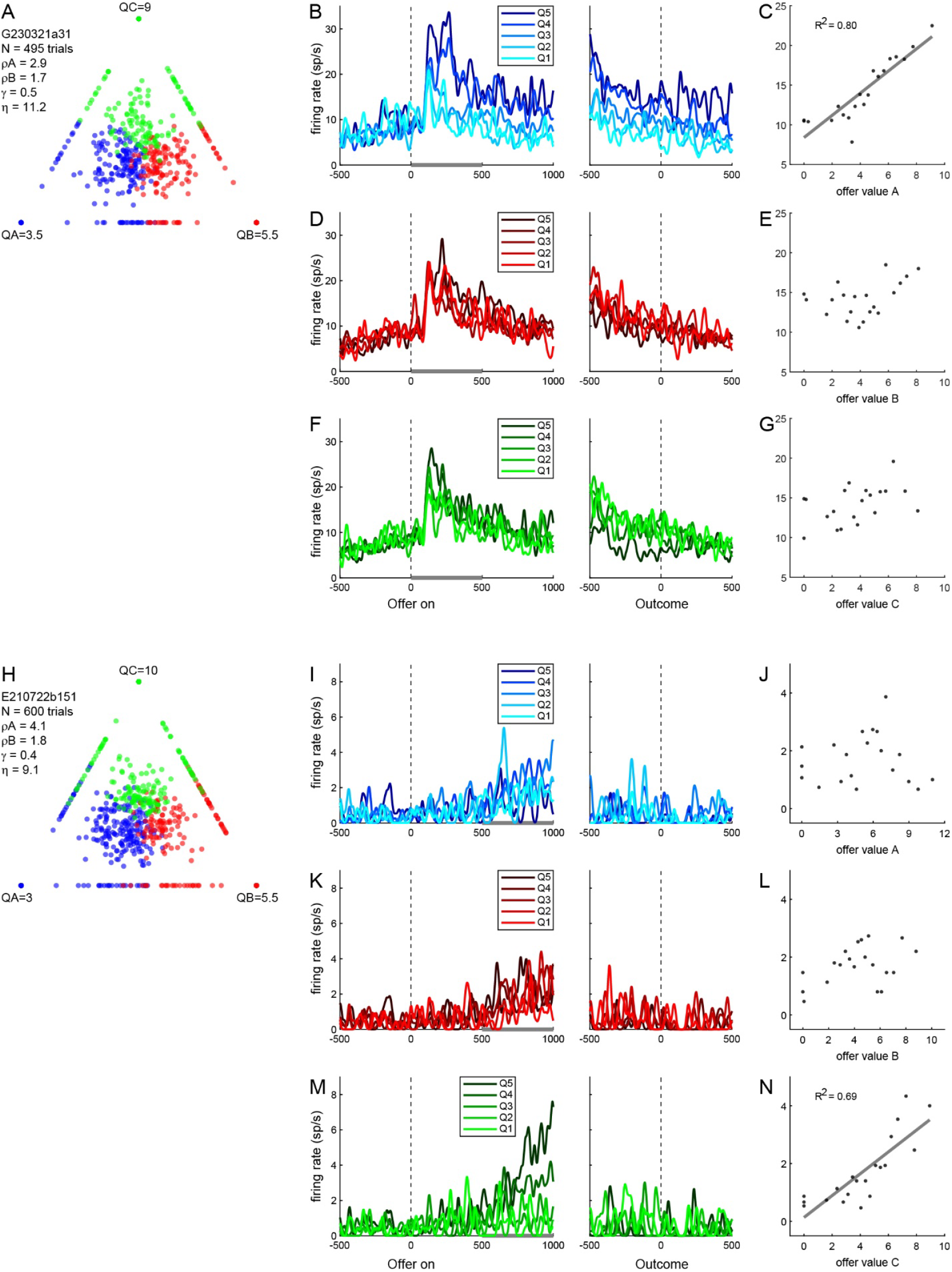
Example neurons encoding individual offer values. **A-G**. Cell encoding the offer value A. Panel A illustrates the behavior. The number of trials and the parameters derived from the logistic regression are indicated. For panel B, trials were divided into 5 quantiles according to the offer value A. For each quantile (Q1 … Q5), we computed the cell activity profile by averaging firing rates across trials (see **Methods**). Starting ∼150 ms after offer onset and until the trial end, the activity of this neuron appeared modulated by the offer value A. Panel C illustrates the activity in the post-offer time window (highlighted in gray in panel B). Here trials were divided into 20 quantiles according to the offer value A. For each quantile, spike counts were averaged over the time window and across trials. The resulting firing rates (y-axis) are plotted here against the offer value A (x-axis). The gray line is from a linear regression and the R^2^ is indicated in the panel. Notably, there was a tight linear relation between firing rates and the encoded variable. In panels DE, the same neuron is examined in relation to variable offer value B. In this case, firing rates don’t appear related to the variable at any time during the trial (panel D). In particular, the regression slope obtained in the post-offer time window (panel E) did not differ significantly from 0. Similar results were obtained when we examined the neuron in relation to variable offer value C (panels FG). **H-N**. Cell encoding the offer value C. Panels JLN illustrate the activity in the late delay time window. The activity of this cell was modulated by the offer value C (panels MN) and did not depend on either the offer value A (panels IJ) or the offer value B (panels KL).

Other neurons appeared to encode variables reflecting the animal’s choices. **Fig.4A-E** illustrates one example cell. In panel B, trials were divided into 5 quantiles according to the *chosen value*. The cell activity was strongly modulated by this variable throughout the offer presentation. In particular, the firing rate recorded in the 500 ms following the offer onset (post-offer time window) showed a tight, approximately linear relation with the *chosen value* (**Fig.4C**). In contrast, the cell activity was effectively independent of the chosen juice (**Fig.4DE**). Another example neuron is illustrated in **Fig.4F-J**. In this case, the cell activity appeared to encode the *chosen value* with a negative slope (negative tuning). A third cell is illustrated in **Fig.4K-O**. The activity of this neuron was strongly correlated with the chosen juice – high when the animal chose juice C and low when the animal chose either juice A or juice B (**Fig.4NO**). The activity of this cell was also mildly correlated with the *chosen value*, which likely reflected the fact that variables *chosen juice C* and *chosen value* were correlated in this session (see **Fig.4K**).

**Figure 4.**
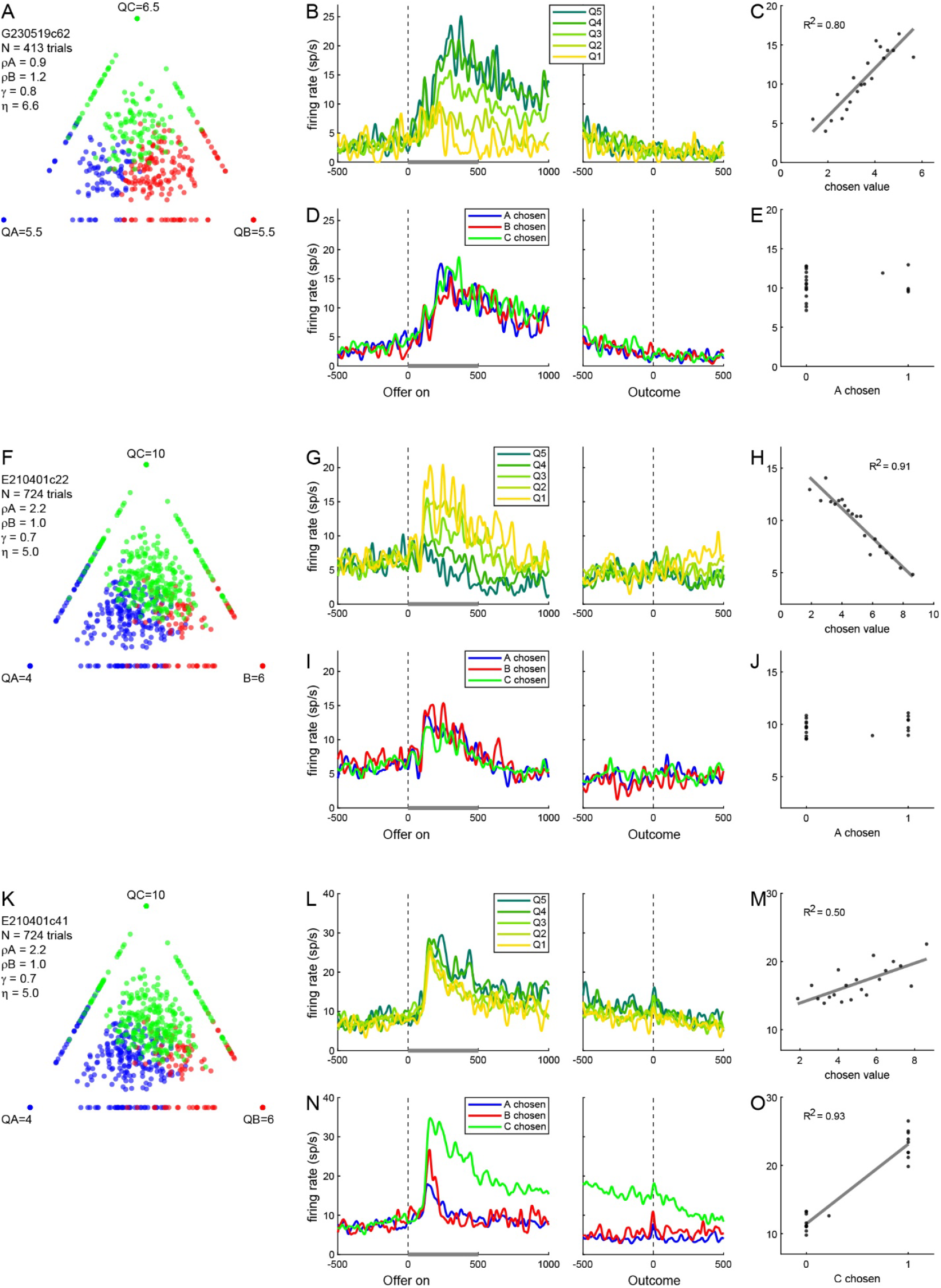
Example neurons reflecting the animal’s choice. **A-E**. Chosen value cell, positive encoding. For panel B, trials were divided into 5 quantiles according to the chosen value and we computed the cell activity profile for each quantile (Q1 … Q5). During offer presentation, the activity of this neuron appeared modulated by the chosen value. Panel C illustrates the activity in the post-offer time window (same format as **Fig.3B**). In panels DE, the same neuron is examined in relation to the chosen juice. Here, trials were divided into three groups depending on whether the animal chose juice A, B, or C (see legend). Firing rates did not appear to reflect this variable. **F-J**. Chosen value cell, negative encoding. Panels HJ illustrate the cell activity in the post-offer time window. This neuron represented the chosen value with negative encoding (panels GH) and did not depend on the chosen juice (panels IJ). **K-O**. Cell encoding the chosen juice C. Panels MO illustrate the cell activity in the post-offer time window. The activity of this neuron was roughly binary – high when the animal chose juice C and low when it chose either juice A or juice B (panels NO). Firing rates were also mildly correlated with the chosen value (panels LM). Most likely, that is because offer values for juice C were higher on average than offer values of the other two juices (see panel K). Note that the two neurons in panels F-J and K-O were recorded in the same session.

Finally, for a sizable group of neurons, firing rates late in the trial were strongly modulated by the outcome – i.e., on whether the chosen juice was delivered or not delivered to the animal. **Fig.5A-E** illustrates one example cell whose activity late in the trial encoded the variable *got juice*. For this cell, the firing rate increased when the animal did not receive the chosen juice (**Fig.5BC**). Importantly, the activity did not depend on what juice the animal had chosen (**Fig.5DE**). A different example neuron is illustrated in **Fig.5FH**. In this case, the cell activity was enhanced only when the monkey chose and received juice A (**Fig.5G**). In other words, this neuron seemed to encode the variable *A received* (**Fig.5H**).

**Figure 5.**
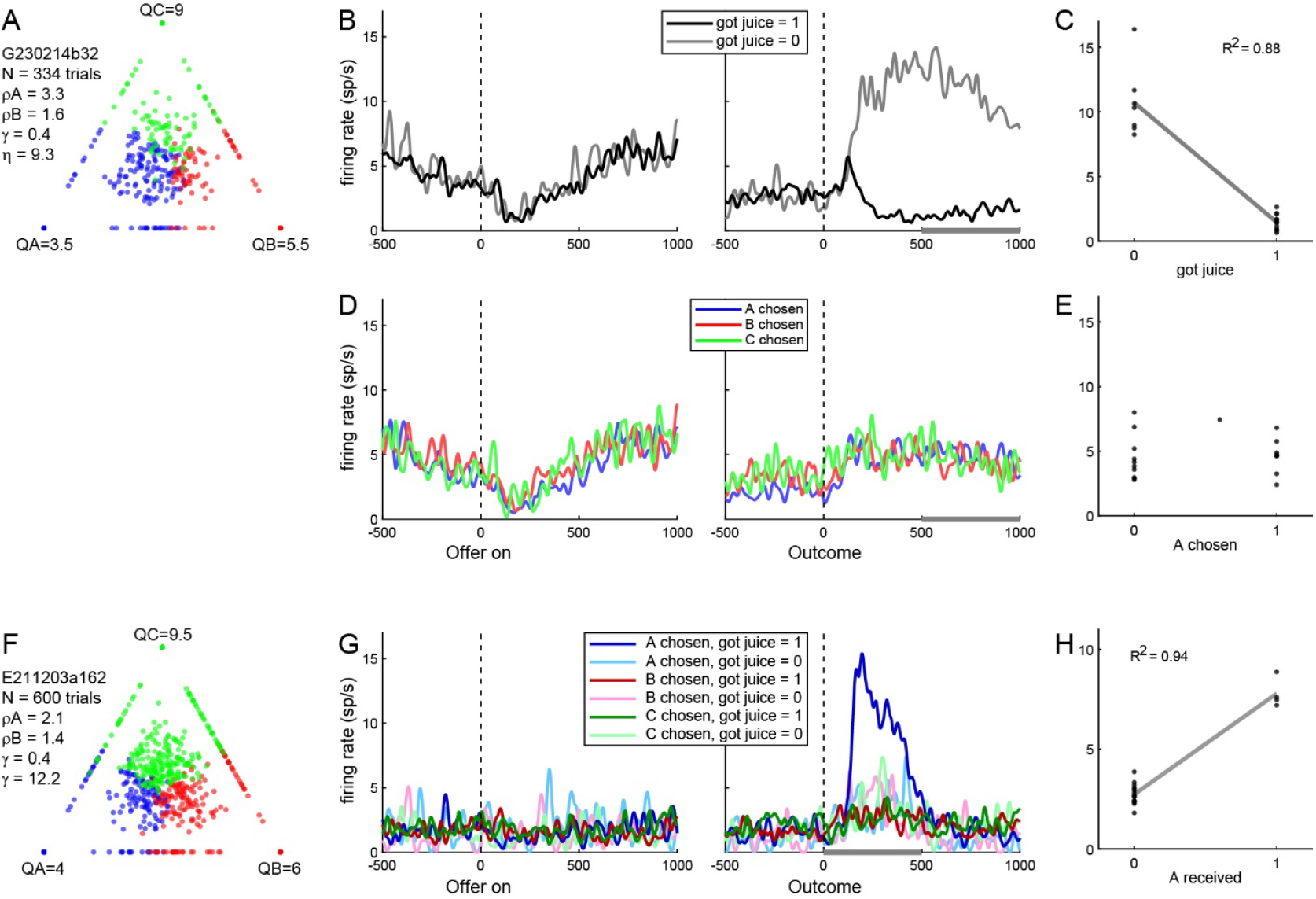
Example neurons encoding the trial outcome. **A-E**. Cell encoding the variable got juice. For panel B, trials were divided into 2 groups according to whether the juice was delivered or not delivered (poor luck) at the end of the trial. Late in the trial, the activity of this neuron was strongly modulated by this variable. In panels DE, the same neuron is examined in relation to the chosen juice. Here, trials were divided into three groups depending on whether the animal chose juice A, B, or C. Firing rates did not appear to reflect this variable. Panels CE illustrate the cell activity in the post-outcome2 time window. **F-H**. Cell encoding the variable juice A received. In panel G, trials were divided in 6 groups according to the chosen juice (A, B, C) and the trial outcome (juice received or not received). After the trial outcome, this cell responded selectively in trials where juice A was chosen and received. In all other circumstances, the cell was essentially not responsive. Panel H illustrates the cell activity in the post-outcome time window.

### Variable selection analysis

For a quantitative analysis of the neuronal population, we adopted a strategy similar to that of previous studies (Padoa-Schioppa and Assad, 2006; Ballesta and Padoa-Schioppa, 2019). We defined six 500 ms time windows aligned with offer onset (go signal), offer offset, and juice delivery (see **Methods**). A neuronal response was defined as the activity of one cell in one time window. We then defined 29 candidate variables that neurons in OFC could conceivably encode (**Table 1; Fig.6**). Some of these variables characterized individual offers. Specifically, for each juice J = A, B, C, we defined variables *offer quantity J, offer probability J, offer expected quantity J, offer value J*, and *offer risk J*. Other variables reflected the animal’s choice (e.g., *chosen quantity, chosen probability, chosen value*, etc.). Among these, we included a spatial variable reflecting the chosen hemifield. Finally, some variables reflected the trial outcome (*got juice, received value*, etc.). We then regressed each response on each variable, as follows.

**Figure 6.**
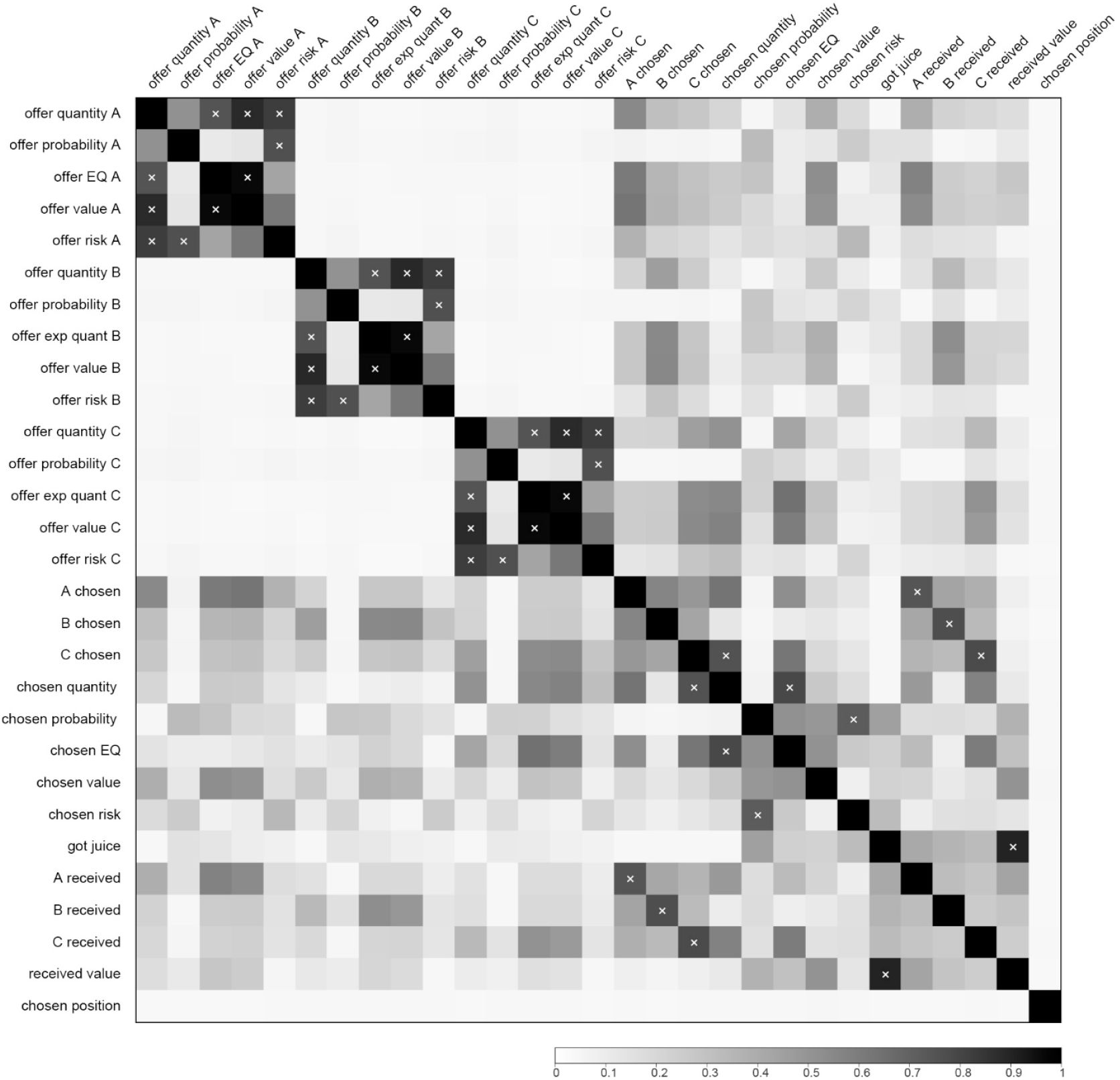
Pairwise correlation between decision variables. The variables examined in the analysis of neuronal activity (**Table 1**) were correlated with each other. For each pair of variables X and Y and for each session, we computed the pairwise correlation r(X, Y) across trials. The gray scale image shown here illustrates the absolute value of the pairwise correlation (|r(X, Y)|) averaged across sessions. Pairs of variables for which the mean correlation was >0.7 are highlighted with a white x symbol.

For each variable, we binned trials in 20 quantiles (each including the same number of trials). For each quantile, we averaged spike counts across trials, and we regressed the resulting firing rates against the variable (see **Methods**). We thus obtained a regression slope and the R^2^. If the slope differed significantly from zero (p<0.01), the variable was said to explain the response. If the variable did not explain the response, we arbitrarily set R^2^ = 0. We repeated this procedure for each neuron, for each time window, and for each variable. Neurons explained by at least one variable in one time window were included in subsequent analyses. We then constructed a population table where rows and columns corresponded to candidate variables and time windows, respectively. Each entry in this table indicated the number of neuronal responses explained by the corresponding variable (**Fig.7A**). Importantly, many of the variables defined in **Table 1** were highly correlated. Consequently, any neuronal response could be explained by >1 variable. For each response, we identified the variable that provided the best explanation (highest R^2^), and we constructed a second population table where entries indicated the number of neuronal responses best explained by the corresponding variable (**Fig.7B**).

**Figure 7.**
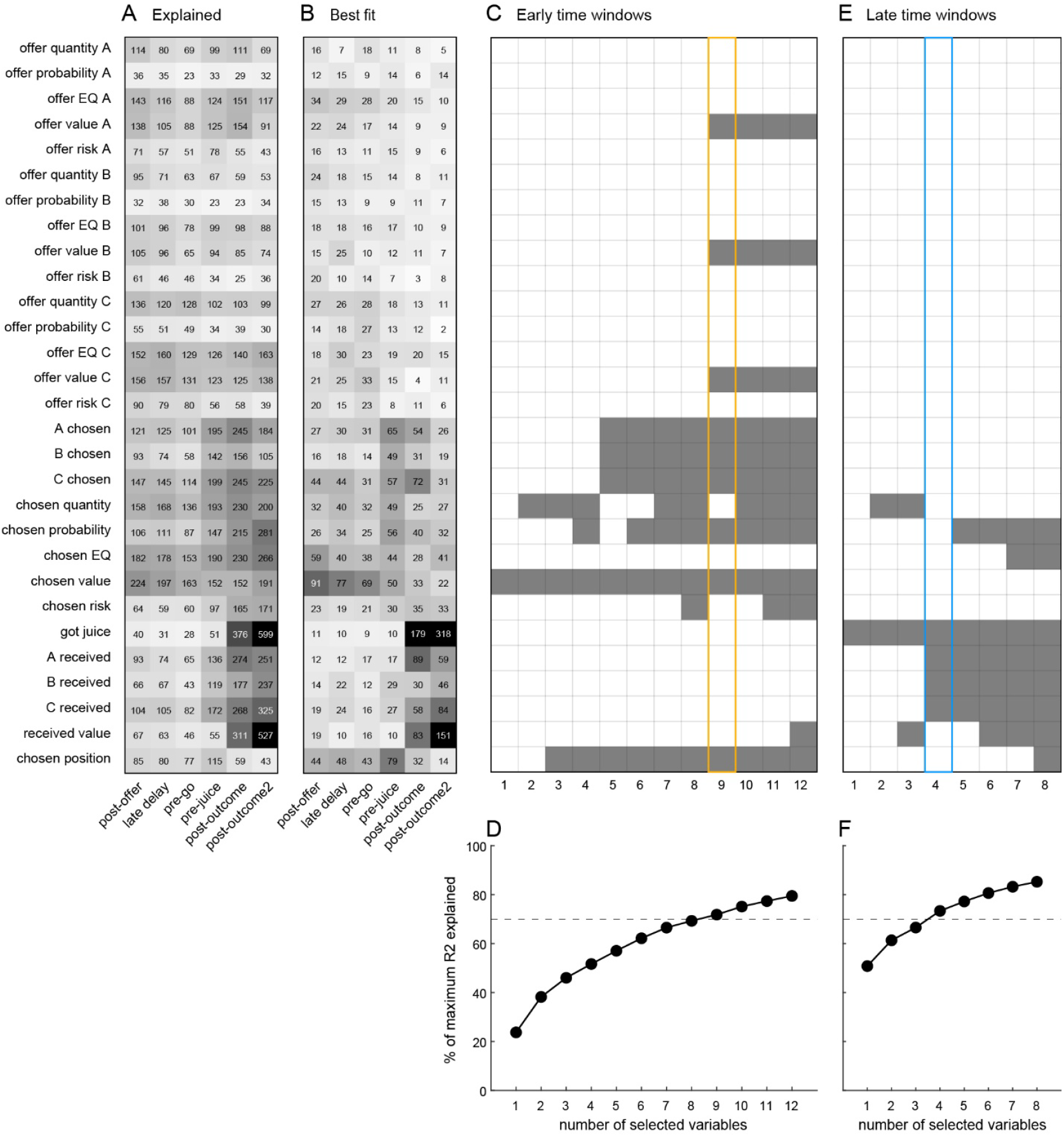
Variable selection analysis. **A**. Explained responses. In this table, rows are candidate variables, columns are time windows, and entries are cell counts. Each entry indicates the number of responses explained by the corresponding variable (non-zero regression slope). Because variables were correlated, many responses were explained by multiple variables, and thus contributed to multiple entries. **B**. Best fit. Here, each response was assigned to the variable providing the best explanation (highest R^2^). Thus each response contributed ≤1 entry. **CD**. Variable selection analysis, early time windows. For *k* = 1, 2, …, we identified the subset of k variables providing the maximum explanatory power (maximum total R^2^). In panel C, gray shades indicate the subset of selected variables (rows) for increasing values of k (columns). Panel D illustrates the explanatory power as a function of k. The dashed line indicates 70% and 100% is the total R^2^ explained by the 29 variables. For *k* = 9, selected variables include 3 *offer value* variables, 3 *chosen juice* variables, *chosen value, chosen probability*, and *chosen hemifield*. **EF**. Variable selection analysis, late time windows. Same format as in panels CD. For k = 4, selected variables include *got juice* and the 3 *juice received* variables.

We aimed to identify a smaller subset of variables that could explain the bulk of the neuronal population. Following previous studies, we conducted a variable selection analysis using a best-subset procedure (exhaustive search). For *k* = 1, 2, 3, … we examined every subset of *k* variables while imposing a consistency constraint (see **Methods**). We then computed the corresponding total R^2^, defined as the sum of the R^2^ accounted for by the *k* variables across the population. The subset providing the largest total R^2^ was identified as the best subset of *k* variables. The procedure was then repeated for increasing values of *k*, until the best subset of *k* variables explained at least 70% of the total R^2^ explained by the full set of 29 variables. Importantly, early time windows (preceding the trial outcome) and late time windows (following the trial outcome) were analyzed separately.

**Fig.7C** illustrates the results of this procedure for early time windows. The single variable with the highest explanatory power, namely *chosen value*, explained 24% of the total R^2^ explained by the full set of 29 variables. Naturally, the explanatory power of the best subset increased with the subset size. For example, for *k* = 5, the best subset – including variables *chosen value, A chosen, B chosen, C chosen*, and *chosen hemifield* – explained 57% of the total R^2^ explained by the 29 variables. The threshold of 70% was crossed for *k* = 9 (**Fig.7E**). In this case, the best subset included variables *offer value A, offer value B, offer value C, A chosen, B chosen, C chosen, chosen value, chosen probability*, and *chosen hemifield*, which we thus identified as the variables encoded by the neuronal population in early time windows.

We repeated this analysis focusing on late time windows (**Fig.7D**). The single variable with the highest explanatory power, namely *got juice*, explained 51% of the total R^2^ explained by the 29 variables. The threshold of 70% was crossed for *k* = 4 (**Fig.7F**). In this case, the best subset included variables *got juice, A received, B received*, and *C received*, which we identified as the variables encoded by the neuronal population in late time windows.

Importantly, the results of the variable selection analysis were quite robust with respect to *k*. The consistency constraint imposed in the procedure (see **Methods**) implied that 3 analogous variables associated with a single juice were either selected together or not selected at all. Inspection of **Fig.7C** (early time windows) illustrates this point for variables *A chosen, B chosen*, and *C chosen* (selected for *k* ≥ 4) and, separately, for variables *offer value A, offer value B*, and *offer value C* (selected for *k* ≥ 9). Notably, once selected, both triplets of variables remained in the best subset even when *k* was increased. The same was true for variables *chosen value* (selected for *k* ≥ 1) and *chosen hemifield* (selected for *k* ≥ 7). A similar situation can be observed in **Fig.7E** (late time windows). Variable *got juice* (selected for *k* ≥ 1) and the triplet of variables *A received, B received, C received* (selected for *k* ≥ 4) all remained in the best subset even when *k* was increased. Hence, although the threshold set at 70% was arbitrary, in a deeper sense the results emerging from this analysis did not substantially depend on it.

On the basis of these results, we classified each response as encoding one of the selected variables. Specifically, each response was assigned to the selected variable providing the largest R^2^. Responses not explained by any selected variable were classified as untuned.

### Neuronal origins of IIA

We next examined properties of neuronal activity in OFC that might effectively instantiate the IIA observed behaviorally. In broad terms, this question pertains to one aspect of menu invariance. Previous work found that OFC activity is invariant to changes in menu – i.e., the activity of neurons associated with one good (X) did not depend on what other good (Y or Z) was offered as an alternative (Padoa-Schioppa and Assad, 2008). Here the question was whether neuronal activity in OFC is invariant to changes in menu size – i.e., whether the activity of neurons associated with one good (X) depended on whether one other good (Y or Z) or two other goods (Y and Z) were offered in alternative. Focusing on early time windows, we considered two aspects of this issue.

First, we examined whether neurons encoded the same variable under binary choices and under trinary choices. For each neuron and for each time window, we examined the two sets of trials separately. We classified the two neuronal responses according to the R^2^ as described above. We then constructed a population table where columns and rows were the variables encoded in binary and trinary choice trials, respectively, and entries were cell counts (**Fig.8A**). Neurons encoding the same variable in binary and trinary choice trials populated the diagonal of this table. Since different variables were encoded by different fractions of neurons, differences in this table were not immediately interpretable. Thus we computed the corresponding table of odds ratios (ORs; **Fig.8B**), in which entries expressed the strength of the association between the variable encoded under binary and that encoded under trinary choices. For each entry, OR = 1 was the chance level; OR < 1 and OR > 1 indicated that the corresponding cell counts in **Fig.8A** were below chance and above chance, respectively. Three aspects of the OR table (**Fig.8B**) are noteworthy. First, most entries in the bottom row and in the rightmost column were significantly below chance (p<0.01; Fisher’s exact test). In other words, few cells were tuned exclusively in binary or trinary choices. Second, all diagonal entries were significantly larger than chance (p<0.01; Fisher’s exact test). In other words, the frequency of neurons encoding the same variable in binary and in trinary choices was consistently higher than chance. Third, a few off-diagonal entries were significantly larger than chance. In principle, this finding might be due to the fact that the classification in binary choices was based on fewer trials (roughly one half) and was thus noisier than that in trinary choices. To confirm this point, we conducted a bootstrap analysis where we randomly divided trials in two sets of unequal numbers (small and large) matching the number of trials for binary and trinary choices, respectively. We repeated this operation N = 100 times and we constructed a contingency table analogous to that in **Fig.8A** (not shown). The corresponding OR table (**Fig.8C**) closely resembled the one obtained for binary/trinary choices (**Fig.8B**). Breslow-Day tests for homogeneity of odds ratios performed for each entry provided statistical confirmation for this point (**Fig.8D**).

**Figure 8.**
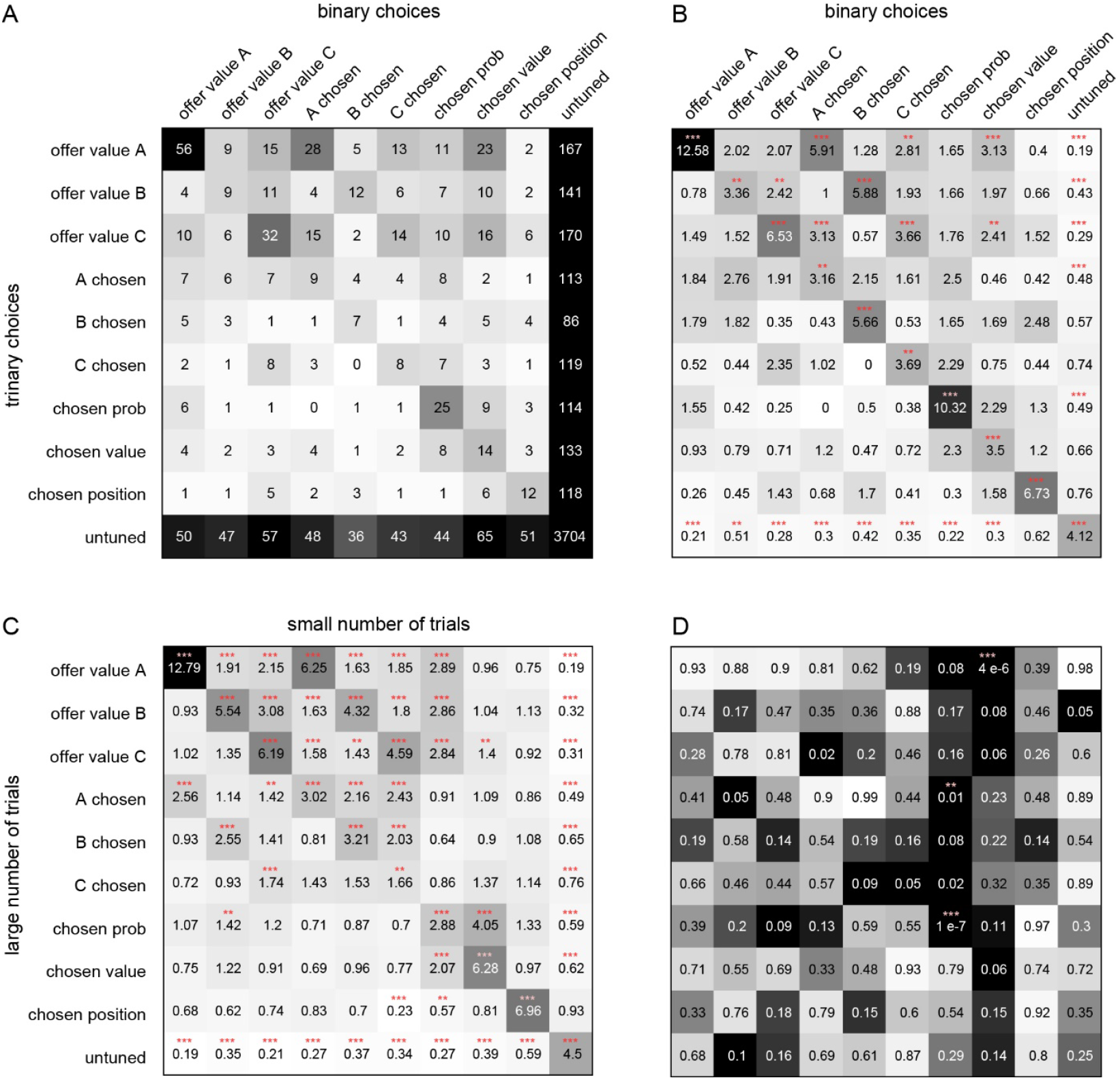
Neuronal origins of IIA. **A**. Variables encoding in binary choices versus trinary choices. Columns and rows are encoded variables in binary and trinary choices, respectively, and entries are cell counts. Shades of gray recapitulate numbers indicated in the table. **B**. Odds ratios (ORs). Entries in this table are ORs. The chance level is OR = 1; OR <1 or OR >1 indicates that the corresponding cell count in panel A was below chance or above chance, respectively. For each entry, red asterisks indicate significant departure from chance (Fisher’s exact test; ^**^ = p<0.01; ^***^ = p<0.001). **C**. Bootstrap analysis, ORs. Equivalent of panel B obtained for bootstrap samples where we randomly divided trials in two unequal sets (large number of trials and small number of trials). Conventions are as in panel B. Notably, the results in panels B and C are qualitatively similar. **D**. Breslow-Day test. Cell counts obtained in panel A and those obtained in the bootstrap analysis were compared using a Breslow-Day test. Numbers and gray shades in this panel are p values. In this analysis, the null hypothesis is that the results obtained with the two manipulations were indistinguishable. Only for 3 of 100 entries in the table reached significance level.

Second, taking for granted that neurons encoded the same variable under binary and under trinary choices, we examined whether variables were encoded with the same activity range (AR) in the two conditions. For each neuronal response, we identified the encoded variable based on all trials. Then we separated binary choices and trinary choices, performed a linear regression against the encoded variable separately for each group of trials, and computed the two ARs (see **Methods**). ARs in binary choices were more broadly distributed and larger on average compared to trinary choices (**Fig.9A-D**). Bootstrap analyses indicated that once the difference in number of trials was accounted for, the ARs measured in binary and trinary choices did not differ (**Fig.9E-L**).

**Figure 9.**
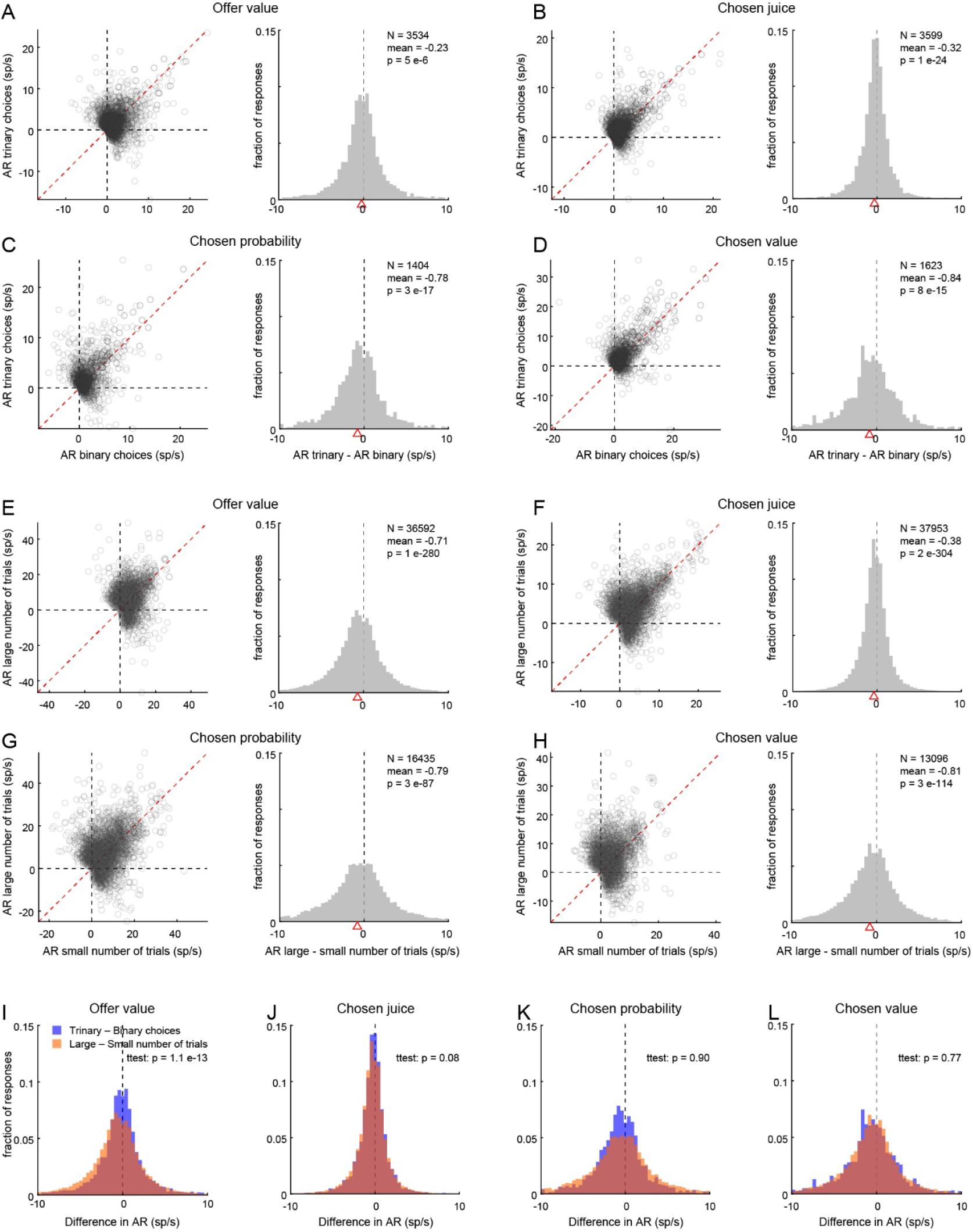
Neuronal origins of IIA, analysis of activity range (AR). **A-D**. Binary choices versus trinary choices. Panel A illustrates the results obtained for offer value responses. For each response, we computed the AR separately in binary and trinary choices. In the scatter plot, each symbol represents one response. Note that data points are more dispersed on the y axis (binary choices) than on the x axis (trinary choices), and that, on average, they lie above the red line (identity). The histogram illustrates the distribution for the difference in AR across the population. The panel indicates the number of responses (N), the mean, and the p value obtained from a t-test. Notably, the mean of the distribution is significantly <0. Similar effects can be observed in panels B, C, and D for chosen juice, chosen probability, and chosen value responses, respectively. **E-H**. Bootstrap analyses. For each variable and each response, trials were divided in two unequal sets (small number of trials; large number of trials). Differences in AR observed for binary vs trinary choices were replicated for sets of small vs large number of trials. **I-L**. Comparing differences in AR. In panel I, the blue and orange histograms are as in panels A and E, respectively. Notably, the yellow histogram is displaced to the left of the red histogram. In other words, splitting trials in unequal sets (bootstrap analysis) over-compensated the effect observed in panel A (which was indeed smaller than those observed in panels B, C, and D). We interpret this observation as due to the fact that binary choices included a relatively large fraction of trials in which the encoded variable (offer value) had value = 0. This was not true for the smaller set constructed for the bootstrap or for other variables. The observed overcompensation is expected if the encoding of offer value is slightly convex.

In summary, our analyses indicated that the encoding of decision variables in OFC is invariant to changes in menu size. If economic decisions are ultimately driven by neuronal activity in OFC, menu size invariance effectively instantiates IIA.

### Offer value cells reflect the subjective nature of economic values

A fundamental question in decision neuroscience is whether value-encoding neurons reflect the subjective nature of economic values. Previous analyses showed that neurons encoding the chosen value in OFC (Padoa-Schioppa and Assad, 2006; Pastor-Bernier et al., 2021) and other areas (Cai and Padoa-Schioppa, 2012; Jezzini and Padoa-Schioppa, 2020, 2024) do. Indeed, session-to-session fluctuations in the activity of these neurons were found to match fluctuations in the relative value of the offered juices. Across sessions, the relation between neuronal and behavioral measures of relative value was statistically indistinguishable from identity (see **Methods**). Furthermore, a study that used different levels of probability showed that session-to-session fluctuations in the activity of chosen value cells in OFC correlated with fluctuations in the risk attitude (Raghuraman and Padoa-Schioppa, 2014). Conversely, previous work did not definitively address this issue for neurons encoding the offer value (Raghuraman and Padoa-Schioppa, 2014). Hence, it remains unclear whether these neurons encode subjective values, as opposed to physical properties of the offers such as the expected quantity of juice. To address this critical issue, we adopted the following strategy. First, we noted that the risk attitude estimated with the logistic analysis fluctuated from session to session for each animal. In principle, such fluctuations could reflect simple measuring errors, but previous work suggests that, in fact, the monkey’s risk attitude can vary over time. Second, we developed an approach do derive a measure for the risk attitude from each offer value response (see below). Third, we compared this neuronal measure (*γ*_*neuron*_) with the behavioral measure obtained from the logistic analysis (*γ*_*behavior*_). More specifically, we examined the relation between the two measures across the population. As detailed in the remainder of this section, the relation between the two measures was statistically indistinguishable from identity.

Our approach to derive a measure of risk attitude from each offer value response was based on the following line of reasoning. If a neuronal response encodes the subjective value of a juice offer, for some parameters *b*_0_ and *b*_1_ the following relation holds true:

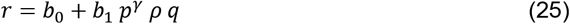

where *r* is the firing rate and the subscript *J* is omitted for brevity. In this light, we divided trials into 6 quantiles according to the offer probability *p*. For each probability quantile, we studied the linear relationship between the firing rate *r* and the offer quantity *q*. That is, we regressed *r* on *q* using the same binning procedure described above (in this case, we used 10 bins). The linear regression was performed for each probability quantile, imposing that the 6 fitted lines have a common intercept (**Fig.10A**). For each regression line (i.e., for each probability quantile), we examined the slope *θ*_*p*_. **Eq.25** implies the following relations:

**Figure 10.**
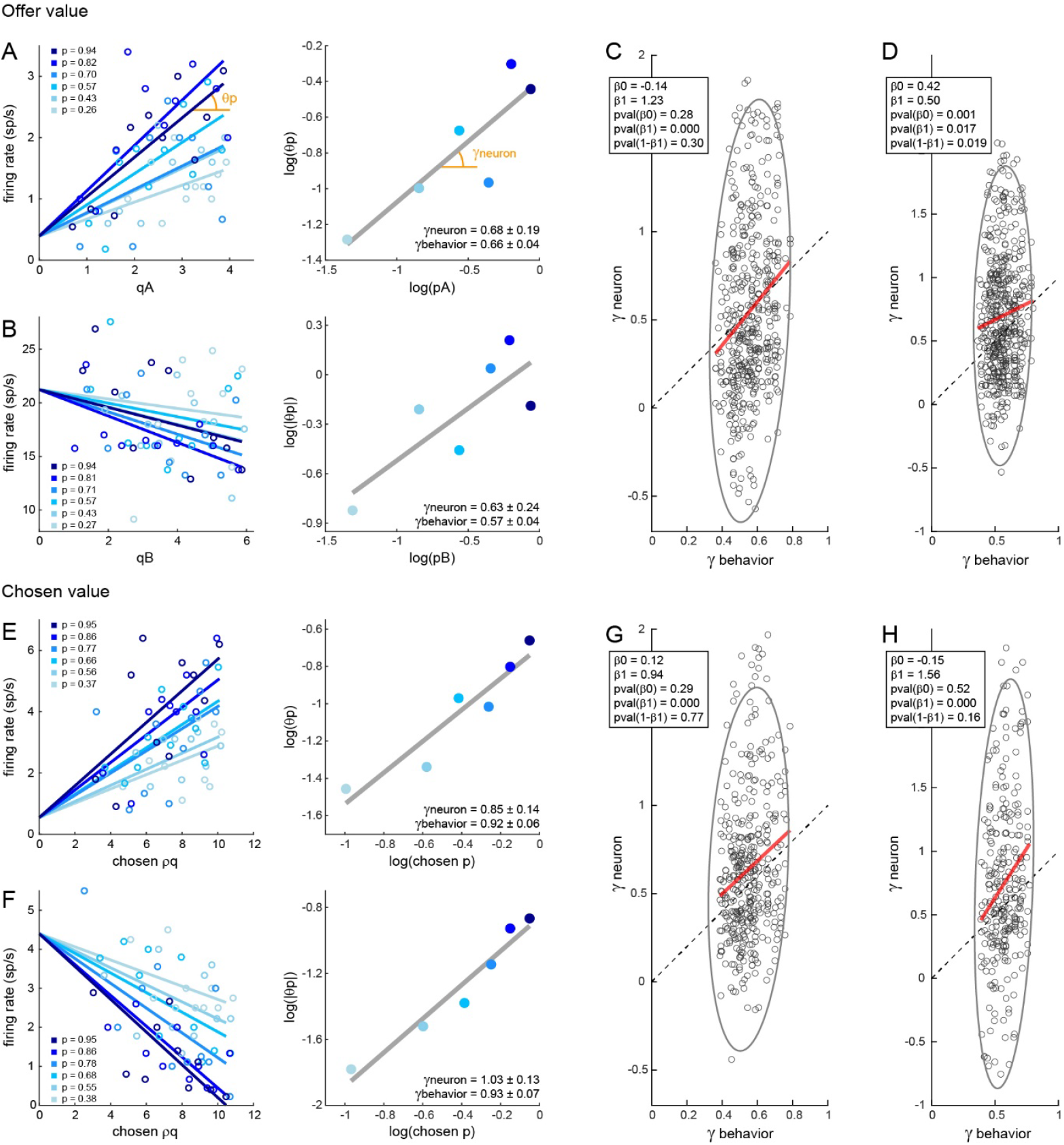
Neuronal and behavioral measures of risk attitude are statistically indistinguishable. **A**. Example offer value response, positive encoding. This neuronal response was classified as encoding the offer value A. Trials were divided into 6 quantiles according to the probability *p*_*A*_. Values of *p* indicated in the figure legend are the means of *p*_*A*_ for each of the 6 quantiles. For each group of trials, we regressed firing rates against the quantity *p*_*A*_, and we examined the regression slope *θ*. We then examined relation between log (*θ*) and log(*p*_*A*_). The slope of the linear fit (± 95% confidence interval) provided the measure for *γ*_*neuron*_. **B**. Example offer value response, negative encoding. **C**. Offer value responses, population analysis. In this plot, x- and y-axis are *γ*_*behavior*_and *γ*_*neuron*_, respectively, and ach data point represents one offer value response (N = 523; positive and negative encoding pooled). The black dashed line indicates identity; the gray ellipse indicates 95% confidence interval for the distribution of data points. The red solid line was derived from linear regression *γ*_*neuron*_ = *β*_0_ + *β*_1_ *γ*_*behavior*_. As indicated in the insert, *β*_0_ was statistically indistinguishable from 0 (p = 0.28); *β*_1_ was significantly different from 0 (p = 3.6 × 10^-8^) and statistically indistinguishable from 1 (p = 0.30). In other words, the relation between the two measures of *γ* was statistically indistinguishable from identity. **D**. Control analysis. Same as panel C, except that the population of responses included in this plot was identified by substituting offer value variables with expected quantity variables in the set of selected variables (see main text). As expected, here *γ*_*neuronal*_ was generally larger than *γ*_*behavioral*_ However, the two measures were significantly correlated across the population – that is, *β*_1_ was significantly >0 (p = 0.017). **EF**. Examples chosen value responses, positive and negative encoding. Same conventions as in panel A. **G**. Chosen value responses, population analysis (N = 423; positive and negative encoding pooled). Again, the relation between *γ*_*neuron*_ and *γ*_*behavior*_was statistically indistinguishable from identity. **H**. Control analysis. Same as panel G, except that the population of responses included in this plot was identified by substituting variable chosen value with variable chosen expected quantity in the set of selected variables. As expected here, *γ*_*neuronal*_ was generally larger than *γ*_*behavioral*_. However, the two measures were significantly correlated across the population – that is, *β*_1_ was significantly >0 (p = 1.24 × 10^-4^).

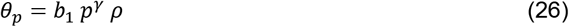

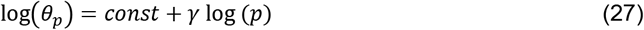

Finally, we regressed log(*θ*_*p*_) on log(*p*). In this regression, each data point represented one probability quantile. The slope of the fitted line provided an estimate for *γ*, referred to as *γ*_*neuron*_.

We applied this analysis to an example response encoding the *offer value A* (**Fig.10A**), from which we obtained *γ*_*neuron*_ = 0.68 ± 0.19 (95% confidence interval derived from the last linear regression). This measure was statistically indistinguishable from that obtained from the logistic analysis, namely *γ*_*behavior*_= 0.66 ± 0.04 (95% confidence interval derived from the logistic analysis).

**Fig.10B** illustrates the procedure for another example response encoding the *offer value B* with a negative slope. Again, we divided trials into 6 probability quantiles. For each probability quantile, we regressed the firing rate against the offered quantity imposing the same intercept for each probability quantile. In this case, *θ*_*p*_ < 0, and we examined the absolute value |*θ*_*p*_|. We regressed log|*θ*_*p*_| on log (*p*), and obtained the regression slope *γ*_*neuron*_ = 0.63 ± 0.24. This measure was statistically indistinguishable from that obtained behaviorally in this session, namely *γ*_*behavior*_= 0.57 ± 0.04.

That the neuronal measures for *γ* obtained for the two responses in **Fig.10AB** were indistinguishable from the corresponding behavioral measures could be completely accidental. However, the risk attitude shown by the monkeys varied from session to session (see **Fig.2D**). We thus examined the relationship between the two measures of *γ* (**Fig.10C**). Unsurprisingly, neuronal measures were generally more dispersed (larger errors of measure). Most importantly, the linear relationship between the two measures computed across the population was statistically indistinguishable from identity (i.e., the intercept was indistinguishable from 0 and the slope was indistinguishable from 1).

Importantly, **Fig.10C** included all neuronal responses classified as encoding an offer value. One concern was that this procedure introduced some circularity because offer value responses were identified for high correlation with offer value variables, which incorporated *γ* (see previous section). Thus for a control analysis, we reclassified each neuronal response in the population while substituting offer value variables (#4, #9, #14 in **Table 1**) with expected quantity variables (#3, #8, #13 in **Table 1**) in the set of selected variables. Naturally, the pools of responses classified as offer value by the two procedures were highly overlapping, but they were not identical. We then repeated the analysis described above for the control pool of responses, and we examined the relationship between *γ*_*neuron*_ and *γ*_*behavior*_. We expected that the control procedure would bias *γ*_*neuron*_ towards 1 (because expected quantity variables effectively set *γ* = 1). However, if most of the responses in the pool indeed encoded the subjective value of the offer, the measures of *γ*_*neuron*_ and *γ*_*behavior*_should remain highly correlated. Our analysis confirmed both these predictions (**Fig.10D**). In summary, these results demonstrated that the activity of offer value cells in OFC reflected the subjective nature of economic values.

We conducted analogous analyses for neuronal responses encoding the *chosen value*. Notably, **Eq.25** held true, but in this case the right-hand side of the equation referred to the chosen option (hence, *ρ* varied from trial to trial). For each response, we divided trials in 6 quantiles according to the chosen probability *p*. For each probability quantile, we regressed firing rates *r* against the chosen *ρ q* imposing that the 6 regression lines have the same intercept. We then examined the 6 slopes *θ*_*p*_. Specifically, we regressed log*θ*_*p*_ on log(*p*), and we obtained the regression slope *γ*_*neuron*_. For responses with negative encoding, we used the same procedures but we rectified *θ*_*p*_. **Fig.10EF** illustrates the results obtained for two example cells with positive and negative encoding. In both cases, the measure obtained for *γ*_*neuron*_ was statistically indistinguishable from the measure of *γ*_*behavior*_obtained from the logistic analysis. Finally, we proceeded with a population analysis of the relation between *γ*_*neuron*_ and *γ*_*behavior*_. First, we classified all neuronal responses based on the set of variables emerging from the variable selection analysis (previous section). The current analysis of the pool of chosen value responses indicated that the relation between *γ*_*neuron*_ and *γ*_*behavior*_was statistically indistinguishable from identity (**Fig.10G**). Again, one concern was that this procedure introduced some circularity. Thus for a control analysis, we reclassified all neuronal responses while substituting the variable chosen value (#22 in **Table 1**) with the variable chosen expected quantity (#21 in **Table 1**) in the set of selected variables. As expected, at the population level, this control procedure biased *γ*_*neuron*_ towards 1. Most importantly, however, the measures of *γ*_*neuron*_ and *γ*_*behavior*_remained highly correlated (**Fig.10H**). In conclusion, these results confirm that the activity of chosen value cells reflected the subjective nature of values.

## DISCUSSION

The choice task adopted here expanded those used in previous studies in three ways: (1) options varied in three dimensions (juice flavor, juice quantity, and probability), (2) quantity and probability varied continuously and were not quantized, and (3) binary and trinary choices were randomly interleaved in each session. This design allowed us to address critical questions left open by earlier work. We reported four primary results. First, our monkeys’ choices were consistent with IIA. Second, neurons in OFC encoded individual offer values, the choice outcome (chosen juice), and the chosen value; these variables generalize those identified under binary choices. Third, the representation of decision variables in OFC was invariant to changes in menu size, which implies IIA. Fourth, the activity of offer value cells reflected the subjective nature of value. We discuss each finding in turn.

IIA is a fundamental assumption of standard economic theory. In essence, IIA holds true if the relative preference between two options X and Y is not affected by introducing a third option Z. Conversely, if the presence of option Z alters the relative preference between X and Y, IIA is violated. Previous work that examined the effects of including a third option in the choice set reported a broad spectrum of results. In humans, numerous studies found multiple forms of IIA violations (Huber et al., 1982; Soltani et al., 2012; Dumbalska et al., 2020). However, other studies did not find systematic violations of IIA (Spektor et al., 2021). Similarly, significant violations of IIA have been observed in non-human species including amoebas (Latty and Beekman, 2011), bees (Shafir et al., 2002), birds (Shafir et al., 2002; Bateson et al., 2003), and primates (Louie et al., 2013; Marini et al., 2023; Marini et al., 2024). Yet, in other studies, animals did not systematically violate IIA (Edwards and Pratt, 2009; Cohen and Santos, 2017; Pastor-Bernier et al., 2017; Parrish et al., 2018; Sánchez-Amaro et al., 2019). The reasons for such a variety of findings are not well understood. One emerging concept is that IIA violations depend on the details of the experimental design. For one, it is critical to distinguish between bona fide IIA violations and analogous – but fundamentally different – perceptual phenomena. In some cases, what appears to be a failure of the choice mechanism might reflect a perceptual illusion (Lea and Ryan, 2015; Parrish et al., 2015; Zhen and Yu, 2016). That aside, IIA violations in humans are more reproducible when offer attributes, such as quantity and probability, are described explicitly (i.e., verbally). Conversely, when the representation of offers is based on experience, IIA violations are less consistent or even reversed (Barron and Erev, 2003; Ert and Lejarraga, 2018; Hadar et al., 2018; Spektor et al., 2019). In other species, where the representation of offers is necessarily experience-based, IIA appears more likely to hold if one of the dimensions defining the offers is categorical as opposed to continuous. For example, this occurs when options are different types of foods (Cohen and Santos, 2017; Sánchez-Amaro et al., 2019) or different tasks (Parrish et al., 2018). In addition, when IIA is violated, the effect size can depend on the deliberation time and on whether that time is internally or externally controlled (Spektor et al., 2021). In summary, the presence or absence of IIA violations cannot easily be linked to a single factor. Our study does not resolve this issue in general. Anecdotally, prior to conducting the present study, we experimented in one animal with a variety of task designs, hoping to elicit systematic violations of IIA. For example, we used options that varied only in quantity and probability (i.e., a single juice flavor), asymmetrically dominated decoys, and asymmetrically dominant phantom decoys. We were unable to replicate classic effects described in humans. Of course, more systematic behavioral experiments should be conducted on a larger cohort. That said, the behavioral results of the present study are noteworthy because we examined a large data set (170-725 trials per session, 248 sessions), because offers in these experiments varied on 3 dimensions – one categorical (juice type) and two continuous (quantity, probability) – and because our analysis examined possible IIA violations in a very broad sense, contrasting choice accuracy in binary versus trinary choices.

An early study examined the OFC of monkeys choosing between different juice flavors offered in variable amounts. It identified distinct neural responses encoding individual offer values, the chosen juice, and the chosen value (Padoa-Schioppa and Assad, 2006). Subsequent work confirmed this result (Kimmel et al., 2020; McGinty and Lupkin, 2023) and extended it to choices under risk (Raghuraman and Padoa-Schioppa, 2014), choices under variable action cost (Cai and Padoa-Schioppa, 2019), choices between juice bundles (Pastor-Bernier et al., 2019), choices under sequential offers (Ballesta and Padoa-Schioppa, 2019; Yun et al., 2020; Shi et al., 2022a), and choices based on decision confidence (Hirokawa et al., 2019). A computationally analogous, but spatial, representation of decision variables was also found in rats (Gore et al., 2023) and mice (Kuwabara et al., 2020; Livi et al., 2025). Our current findings generalize this core result in a critical direction – i.e., choices between multiple options. That neurons in OFC essentially represent the same decision variables in ternary as in binary choices is consistent with the idea of decision mechanisms that naturally scale to multinary choices (Furman and Wang, 2008; Wang, 2012; Battista et al., 2025). In addition to neurons encoding individual offer values, the choice outcome, and the chosen value, we found cells encoding the chosen probability and the chosen hemifield (or chosen action). The latter finding is somewhat surprising and seemingly at odds with previous results (Padoa-Schioppa and Assad, 2006; Grattan and Glimcher, 2014). One possibility is that the more complex task design induced a partly-spatial representation. Future studies should examine this hypothesis.

Leveraging the fact that binary and trinary choices were randomly interleaved in each session, we found that individual neurons encoded the same variable with the same activity range in binary and in trinary choices. In other words, the representation of decision variables was invariant to changes in menu size. This observation is especially noteworthy for neurons encoding the offer values because one could reasonably hypothesize that the activity of these neurons does, in fact, depend on the menu size. Indeed, consider the activity of a cell encoding the offer value X recorded while offer X is presented to the animal. Under binary choices, only one other offer competes for visual attention and cognitive resources; under trinary choices, two other offers compete for cognitive resources. In this light, one might predict that offer value X signals recorded under trinary choices are somewhat degraded compared to those recorded under binary choices. That this was not the case is, from our perspective, quite remarkable. It resonates with findings from an earlier study where we interleaved choices under simultaneous and sequential offers: surprisingly, offer value signals recorded under simultaneous offers (when two offers competed for attentional resources) were stronger than those recorded under sequential offers (when options were evaluated one at the time) (Shi et al., 2022b). Taken together, the results of the two studies suggest that the strength of the value representation is robust and not simply governed by sensory constraints. If economic decisions are indeed formed within OFC, invariance to changes in menu size effectively instantiates IIA. Importantly, using different task designs, it might be possible to elicit systematic violations of IIA. In such conditions, we would expect the neuronal representation in OFC to vary with the menu size.

Finally, we addressed a long-standing issue – i.e., whether the representation of offer values in OFC reflects the subjective nature of value. The alternative hypothesis is that neurons nominally encoding the offer value actually represent objective properties of the offers such as the expected quantity. Earlier work addressed this fundamental question for chosen value cells. Specifically, the spiking activity of these cells provided neuronal measures for the relative value of two juices (Padoa-Schioppa and Assad, 2006; Pastor-Bernier et al., 2021) and for the risk attitude (Raghuraman and Padoa-Schioppa, 2014). These measures fluctuated from session to session and were statistically indistinguishable from the corresponding behavioral measures. In contrast, previous work did not provide conclusive evidence for offer value cells. The main difficulty in nailing this issue follows from the fact that offer value cells undergo range adaptation (Padoa-Schioppa, 2009). Consequently, session-to-session fluctuations in relative values are not matched in the activity of these neurons. An alternative approach is to examine whether fluctuations in the activity of offer value cells matches behavioral fluctuations in the risk attitude (Raghuraman and Padoa-Schioppa, 2014; Lak and Stauffer, 2015). However, an earlier study that attempted this approach lacked the necessary statistical power (Raghuraman and Padoa-Schioppa, 2014). Leveraging the fact that quantities and probabilities varied continuously, here we were able to settle this fundamental issue. In essence, neuronal and behavioral measures of risk attitude co-varied across sessions and their relation was indistinguishable from identity. This finding proves that offer value signals in OFC reflect the subjective nature of value.

## Acknowledgements

We thank members of the Padoa-Schioppa lab for helpful discussions. This work was supported by the National Institutes of Health (grant number R01-DA032758).

## Notes

**Conflict of interest:** None

### Competing Interest Statement

The authors have declared no competing interest.

